# Lung cancer-enriched p53 mutants occupy canonical p53 target genes without activating transcription, revealing a distinct loss-of-function behavior

**DOI:** 10.64898/2026.02.14.705936

**Authors:** Mason A. Tracewell, Hailey N. Shankle, Samantha M. Barnada, Khushali S. Vyas, Kevin M. Kim, Theodhora Qyshkollari, Jonathan E. Karlin, Julie A. Barta, Steven B. McMahon

## Abstract

Lung cancer is the most common cause of cancer-related death in the U.S. and globally. Cigarette smoking remains the leading risk factor for lung cancer, in part by inducing loss-of-function mutations in tumor suppressor genes, including *TP53*. While most cancers share a set of common “hotspot” mutations in p53, lung cancer exhibits an additional, distinct cluster of hotspot mutations. This cluster is typified by the missense mutations *TP53*:p.V157F and *TP53*:p.R158L. While canonical hotspot mutations cause broad misfolding of p53 or eliminate specific DNA contact residues, mechanistic studies of the lung cancer mutants reported here demonstrate that they retain the ability to bind the same genomic sites as wild-type p53. Despite actively binding to traditional p53 target genes, the lung cancer mutants are defective in activating transcription. To our knowledge, this represents the first demonstration of functional inactivation of the p53 tumor suppressor at a point after DNA binding, but prior to target gene activation. Relevant to the sequential inactivation of each p53 allele during cancer progression, the lung cancer mutants block the activity of a wild-type p53 allele when co-expressed in a dominant negative manner. Identification of this loss-of-function mechanism has key implications for therapeutic strategies aimed at restoring p53 function in lung cancer.

## Introduction

Functional inactivation of the p53 tumor suppressor protein is an essential step in the progression of most human cancers [1–3]. Distinct from most tumor suppressors, inactivation of p53 typically involves retention of the entire *TP53* gene in cancer cells [4, 5]. Instead of gene deletion, single amino acid alterations in the p53 protein are introduced as a consequence of point mutations in the underlying locus [6]. This results in tumor cells producing a full-length p53 protein that is non-functional due to the inactivating properties of a single missense mutation [7].

As a sequence-specific transcription factor, wild-type p53 function relies on a highly structured central domain in which several amino acids make direct hydrogen bonds with DNA [8–10]. Many of the DNA contact residues in p53 are hotspots for mutation in cancer, presumably because alterations at these sites prevent the protein from binding to DNA [10]. Additional mutational hotspots in p53 occur at amino acids that trigger protein misfolding, leading to a loss of DNA binding by p53 and a defect in the transcription of target genes related to key cellular processes such as cell cycle control, DNA damage repair, and apoptosis [11–13]. These observations have led to a decades-long search for strategies to “correct” the function of the mutant p53 protein in human cancer [14]. At the biochemical level, these reactivation strategies have focused on refolding p53 into a conformation that restores sequence-specific DNA binding, with numerous clinical trials based on this strategy [15, 16].

We report here that human lung cancer contains a set of missense mutations in the p53 DNA-binding domain that debilitate p53 function by a distinct mechanism. These lung cancer mutations do not impair the ability of p53 to bind its normal sites in the genome and would not respond to current reactivation strategies. Instead, these lung cancer-enriched mutations impair the ability of p53 to induce the transcription of downstream target genes in a non-productive DNA binding manner. The identification of a unique biochemical mode of p53 inactivation has broad implications for the efficacy of reactivation strategies as a therapeutic approach in human cancer.

## Materials and Methods

### Cell Culture

Human NCI-H460 (ATCC HTB-177), NCI-H1299 (ATCC CRL-5083), and NCI-H2087 (ATCC CRL-5922) cells were purchased from the American Type Culture Collection (ATCC, Manassas, VA). Following the receipt of the cell lines, aliquots of passages 2–5 were frozen in liquid nitrogen. New aliquots were thawed every 4–6 months for use in experiments. NCI-H460 and NCI-H2087, cells were maintained in RPMI 1640 (Corning) supplemented with 10% Benchmark fetal bovine serum (100-106, Gemini) or 10% tetracycline (TET) negative fetal bovine serum (100-800, GeminiBio) for the TET-inducible system, 1X Glutamine (100X, A2916801, Gibco) under 37°C and 5% CO_2_ conditions. NCI-H1299 cells were maintained in Dulbecco’s Modified Eagle Medium (Corning) supplemented with 10% tetracycline negative fetal bovine serum (100-800, GeminiBio), 1X Glutamine (100X, A2916801, Gibco) under 37°C and 5% CO_2_ conditions. Cells were passaged twice weekly, and *Mycoplasma* contamination was monitored by a PCR detection kit (30-1012K, ATCC).

Tetracycline-inducible p53-expressing stable cell lines were generated in NCI-H1299 and NCI-H460 cells via lentiviral infection using ViraPower HiPerform Lentiviral Expression System (Invitrogen) [17]. p53 expression vectors were generated in pLenti6.3/V5-DEST cloning vectors (Invitrogen). Expression vectors encoding for Val158 to F (V157F) and Arg158 to L (R158L) p53 mutants (V157F; R158L) were generated using the QuikChange II Site-Directed Mutagenesis kit (Agilent). Expression of wild-type (WT) p53, V157F p53, and R158L p53 were induced with tetracycline (T7660, Millipore Sigma).

### Cellular Images

Cellular images were captured by the EVOS XL Core Imaging System (AMEX1000, Invitrogen by Thermo Fisher Scientific).

### Camptothecin (CPT) Treatment

Camptothecin (J62523.03, Thermo Fisher Scientific) was used at 2μM to investigate p53 response to DNA damage. For TET+CPT conditions, cells were treated with TET for 16 hours to allow for induction of p53, and then CPT was added for 6, 24, or 48 hours.

### Immunoblotting

Cells were harvested and lysed in a Nonidet P-40-based whole-cell lysis buffer supplemented with a protease inhibitor cocktail (P8340, Sigma-Aldrich). Lysate concentration was determined using bicinchoninic acid (BCA) assay and analyzed by SDS–PAGE using antibodies against p53 (sc-126, Santa Cruz Biotechnology), GAPDH (#5174, Cell Signaling Technology), and Caspase-3 (#9662, Cell Signaling Technology). All the antibodies were used 1:1000 in TBS-T.

### CellTiter-Glo 2.0 Cell Viability Assay

The CellTiter-Glo 2.0 cell viability assay (G9242, Promega) assessed cell viability by quantifying the amount of ATP, indicating metabolically active cells. 10-000-25,000 cells were plated per well into a 96-well plate (781971, BrandTech) and allowed to adhere overnight. Per experiment, cells were treated with the indicated treatments and time respectively. The media was removed, and 100μL of fresh media was added per well. 100μL per well of CellTiter-Glo 2.0 reagent was added, and the contents were mixed by a shaker. The plate was incubated at room temperature for 10 minutes. Luminescence was determined using a PolarStar Optima plate reader (BMG LabTech) at room temperature.

### Cell Quantification

Cells were collected with Accutase (07922, Innovative Cell Technologies) and spun down at 1300rpm for 5 minutes. Cells were resuspended with media and 25μL per sample was diluted 1:1 with trypan blue (2680430, Invitrogen). Cells were quantified using Countess 3 (Invitrogen).

### Cell Cycle Analysis by Flow Cytometry

Cells were collected with Accutase and spun down at 1300rpm for 5 minutes. Cells were washed two times with phosphate buffer saline (PBS) (10010-023, Gibco), and then resuspended in PBS. 500,000 cells were placed in a 15mL conical tube with a total volume of 300μL PBS. Cells were fixed by adding 200 proof ethanol dropwise, while vortexing the cells to prevent clumping, to a final concentration of 70% ethanol. Cells were incubated on ice for 30 minutes with vortexing every 5 minutes. Cells were spun down at 1600rpm for 5 minutes and supernatant was aspirated. Cells were resuspended in 50 μL RNAse solution (5μg RNAse in 50μL PBS, R1253, Thermo Fisher Scientific) and 100μL propidium iodide (PI) staining solution (556463, BD Biosciences). Cells were stained for 15 minutes in the dark at room temperature. 350μL of PBS was added to the cell suspension, and then the solution was pipetted into a FACS tube with a cell strainer (352235, Falcon). The data was collected by flow cytometry on a CytoFLEX (Beckman Coulter) using CytExpert (v1.2 Beckman Coulter). Data was analyzed with FlowJo (10.10.0) to perform the cell cycle analysis. Reported data are from a population of 30,000 cells.

### Apoptosis Analysis by Flow Cytometry

Apoptosis was measured using the FITC annexin V apoptosis detection kit (556547, BD Pharmingen) following manufacturer’s instructions. Cells were collected with Accutase and spun down at 1300rpm for 5 minutes. Cells were washed two times with PBS, and then resuspended in 1X binding buffer. 5μL of FITC annexin V and 5μL of PI staining solution was added to 100μL of cell suspension containing 100,000 cells. Samples were incubated at room temperature for 15 minutes in the dark. 400μL of 1X binding buffer was added to each sample. Samples were pipetted into a FACS tube with a cell strainer. The data was collected by flow cytometry on a Cytoflex using Cytexpert. The data was analyzed with FlowJo, and reported data are from a population of 30,000 cells.

### Click-it Reaction

H2087 cells were collected with trypsin (25053CI, Corning) and plated on chamber slides (80427, Ibidi) to adhere for 24 hours at a confluency of ∼70%. Cells were labelled with media containing 5μM 5-ethynyl-2′-deoxyuridine (EdU, 7207, Tocris) for 1.5 hours, followed by 30 minutes of regular media. Cells were washed with PBS and then fixed for 15 minutes with 3.7% formaldehyde (252549, Sigma-Aldrich). Cells were washed with PBS and then permeabilized with 0.25% Triton X-100 (21568-0010, Acros Organics) for 10 minutes. Cells were washed and blocked with 1% BSA (BP1605-100, Thermo Fisher Scientific) for 10 minutes. Click-it reaction was performed for 30 minutes in the dark using 0.1M copper sulfate (C1297, Sigma-Aldrich), 1mM Biotin Azide (900891, Sigma-Aldrich), 0.1M Ascorbic Acid (255564, Sigma-Aldrich) and 0.1x PBS, and then the cells were washed with PBS. DNA biotinylation enabled proximity ligation assay.

### Proximity Ligation Assay

Performed proximity ligation assay (PLA) following manufacturer’s protocol for Duolink In Situ Detection Reagents (DUO92008, Sigma-Aldrich). Cells were blocked for 10 minutes with 1x western blocking solution (11921673001, Roche) and 1.5% normal donkey serum (017-000-121, Jackson ImmunoResearch). The following primary antibodies (one rabbit and one mouse per condition) were added at a 1:1000 dilution: rabbit IGG (NI01, EMD Millipore), Anti-Biotin (rabbit, AB53494, Abcam), or p53 (mouse, sc-126, Santa Cruz Biotechnology). The samples were placed in a humidity chamber at 4°C overnight and remained in the humidity chamber until mounting was completed.

The samples were washed with PBS for 15 minutes and blocked briefly with 1x western blocking solution. The samples were incubated with PLA probes diluted in 1x western blocking solution for 1 hour at 37°C. The samples were washed with Buffer A, and the ligation step was performed for 30 minutes at 37°C. The samples were washed with Buffer A, and the amplification step was performed for 1 hour and 40 minutes at 37°C. The slides were washed with PBS and stored in mounting medium with DAPI (50011, Ibidi). The slides were stored at 4°C until imaging using the DeltaVision Ultra widefield microscope system.

### Luciferase Reporter Assay

The pGL3 firefly luciferase reporter plasmids made for p21 (GAACATGTCCcAACATGTTg) and BAX (GGGCAgGCCCCGGGCTTGCTg) response elements have been described previously [18, 19]. Cells were transfected with 1μg of firefly luciferase reporter plasmid (p21 or BAX), 250ng of p53 expression vectors (WT, V157F, R158L, or EV), and 200ng of pRL-TK Renilla luciferase control reporter vector (E2241, Promega) using Lipofectamine 3000 (L3000015, Invitrogen) according to manufacturer’s instructions. Dual-luciferase reporter assay system kit (E1910, Promega) was used for data collection. Cells were harvested 24 hours post-transfection with passive lysis buffer. Luminescence of samples was detected by a GloMax® Explorer (3.2.3, Promega) and recorded in relative light units (RLU).

### RNA-sequencing

H1299 cells containing a TET-inducible plasmid for WT, V157F, or R158L p53 were treated with TET or TET+CPT as described above and collected. Total RNA was extracted using the RNeasy Plus Mini Kit (74134, Qiagen) following the manufacturer’s protocol. RNA was quantified, and the RNA integrity number (RIN) was verified using an RNA ScreenTape® for Tapestation 4150 (v4.1.1, Agilent). All samples used for library preparation and subsequent sequencing had a RIN value above 8.0. Directional RNA libraries were prepared using 1μg of total RNA as input and using NEBNext^®^ Poly(A) mRNA Magnetic Isolation Module (E7490L, New England Biolabs), NEBNext^®^ Ultra^TM^ II Directional RNA Library Prep Kit for Illumina^®^ (E7760L, New England Biolabs), and NEBNext^®^ Multiplex Oligos Dual Index Primers for Illumina (E7600S and E7780S, New England Biolabs) according to the manufacturer’s instructions. Libraries were sequenced on a NextSeq 2000 (Illumina) generating single-end 150 bp reads.

### RNA-sequencing Analysis

FastQC (https://github.com/s-andrews/FastQC) was used for quality control of all raw fastq files and adapters were removed using TrimGalore! (https://github.com/FelixKrueger/TrimGalore). Reads mapping to each gene in each sample were quantified to abundance using Kallisto [20]. Abundance was then converted to raw gene count via DESeq2 [21]. The raw gene count for each sample was used to determine differential gene expression (p-value < 0.05; FDR <5%). Abundance was used to determine transcript per million for data visualization. All statistical analyses were performed using Kallisto v0.50.0, DESeq2 v1.40.2 [21], R v4.2.3 and Prism 1v0.1.1.

### Reverse Transcription-Quantitative Polymerase Chain Reaction (RT-qPCR) Analysis

Cells were collected and total RNA was extracted using the RNeasy Plus Mini Kit following manufacture’s protocol. cDNA was created from 2μg of RNA using the High-Capacity cDNA Reverse Transcription Kit (4368814, Thermo Fisher Scientific) following manufacturer’s instructions. RT-qPCR was performed on the cDNA using the Fast SYBR^TM^ Green Master Mix (4385612, Thermo Fisher Scientific) following manufacturer’s instructions on the Step One Plus Real Time-PCR system (v2.3, Applied Biosystems). mRNA levels between samples were normalized to *GAPDH* transcript levels. Primer sequences are listed in Supplemental Table 1.

### Chromatin Immunoprecipitation (ChIP)

ChIP was performed as previously described [22] with minor alterations. Approximately 15 million cells were cross-linked with 1% formaldehyde for 10 minutes rocking at room temperature, and then quenched with 125mM glycine (BP381, Fisher Scientific) for 5 minutes. The cells were washed twice with 1X PBS. Chromatin was extracted from the fixed cells as described in [23]. The fixed cell pellet was lysed with ChIP lysis buffer 1 (50 mM Hepes-KOH, pH 7.5; 140 mM NaCl; 1mM EDTA; 10% Glycerol; 0.5% Igepal CA-630; 0.25% Triton X-100), rocked for 10 minutes, and then spun down at 2000xRCF for 5 minutes. The supernatant was aspirated, and the pellet was lysed with ChIP lysis buffer 2 (10 mM Tris-HCL, pH8.0; 200 mM NaCl; 1 mM EDTA; 0.5 mM EGTA), rocked for 5 minutes, and then spun down at 2000xRCF for 5 minutes. The supernatant was aspirated, and the nuclei pellet was lysed with ChIP lysis buffer 3 (10 mM Tris-HCl, pH 8; 100 mM NaCl; 1 mM EDTA; 0.5 mM EGTA; 0.1% Na-Deoxycholate; 0.5% N-lauroylsarcosine). Chromatin was sheared to an average base-pair length of 100-300 using the Q800R2 Sonicator (QSonica) kept at 4°C with circulating chiller (4905-110, QSonica), with 50% amplitude, and a pulse of 20 seconds on and 20 seconds off, for 20 minutes of sonication on time. 80μL TE with 1% SDS was added to 20μL for each sonicated sample. 2μg/μL RNase and 2μg/μL Proteinase K (25530-015, Thermo Fisher Scientific) were added to each sample and incubated for 1 hour at 65°C. Samples were purified with the Monarch® PCR & DNA Cleanup Kit (T1030L, New England Biolabs) following manufacture’s protocol. The chromatin shearing efficiency of the samples was checked by D1000 ScreenTape® Tapestation 4150 (v4.1.1, Agilent).

The remainder of the sonicated samples kept on ice, had Triton X-100 (21568-0010, Thermo Fisher Scientific) added to a final concentration of 1%. The samples were centrifuged at 20,000xRCF for 10 minutes at 4°C. The supernatant was collected, and 2% input samples were collected and stored at -20°C. The remaining sheared chromatin was diluted 1:3 in ChIP lysis buffer 3 with 1% Triton X-100. The sheared chromatin was incubated with 15μg antibodies (p53, AHO0152, Thermo Fisher Scientific; mIGG, sc-2025, Santa Cruz Biotechnology) and Dynabeads^TM^ Protein G beads (10004D, Thermo Fisher Scientific) rotating overnight at 4°C. The beads were washed three times with ChIP wash buffer (0.1% SDS; 1% Triton X-100; 10mM EDTA; 150mM NaCl; 20mM Tris-HCl pH 8.0). The beads were washed a final time with final ChIP wash buffer (0.1% SDS; 1% Triton X-100; 10mM EDTA; 500mM NaCl; 20mM Tris-HCl pH 8.0). The beads were resuspended in 1X TE containing 1% SDS and incubated at 65°C for 10 min, and this was repeated twice to elute all the immunocomplexes. The inputs were diluted to the same volume as the ChIP samples, and the inputs and ChIP samples were incubated overnight at 65°C to reverse cross-linking. The samples were digested with Proteinase K (0.5μg/μL) for 1 hour at 65°C, and then the DNA was purified using the ChIP DNA Clean & Concentrator^TM^ (D5205, Zymo Research).

### ChIP-qPCR

After the samples were purified by the ChIP DNA Clean & Concentrator^TM^ kit, the samples were resuspended with water to a total volume of 100μL. Samples were analyzed by qPCR with 2μL of each ChIP sample per well in triplicate for each ChIP-qPCR primer, ChIP-qPCR primer sequences are found in Supplemental Table 2.

### ChIP-sequencing

After the samples were purified by the ChIP DNA Clean & Concentrator^TM^ kit, the samples were quantified with 1X dsDNA HS Assay Kit (Q33231, Thermo Fisher Scientific) on the Qubit 4 (Thermo Fisher Scientific). Barcoded libraries were prepared with NEBNext^®^ Ultra^TM^ II DNA Library Prep Kit for Illumina^®^ using NEBNext^®^ Multiplex Oligos for Illumina^®^ (Dual Index Primers Set 1) (E7600S, New England BioLabs). Libraries were sequenced on a NextSeq 2000 (Illumina) generating single-end ∼138 bp reads.

### ChIP-sequencing Analysis

FastQC (https://github.com/s-andrews/FastQC) was used for quality control of all raw fastq files and adapters were removed using TrimGalore! (https://github.com/FelixKrueger/TrimGalore). The sequences were then aligned to human reference genome hg19 using the Burrows-Wheeler Alignment tool with the MEM algorithm [24]. Aligned reads were filtered with a mapping quality greater than 10 (MAPQ >10) and PCR duplicates were removed. Peaks were called using MACS2 [25] with default parameters and FDR < 5%. All statistical analyses were performed using BEDTools [26], deepTools [27], R v4.2.3, and Prism 1v0.1.1. ChIP peaks were visualized in Integrative Genomics Viewer [28] on genome build hg19. Only one representative ChIP replicate was used for data visualization purposes.

### Publicly Accessible ChIP-sequencing

For the publicly accessible data used, FASTQ files for each sample were downloaded from GEO: GSE238181 [35] and GSE59176 [36]. FastQC (https://github.com/s-andrews/FastQC) was used for quality control of all raw fastq files and adapters were removed using TrimGalore! (https://github.com/FelixKrueger/TrimGalore). The sequences were then aligned to human reference genome hg19 using the Burrows-Wheeler Alignment tool with the MEM algorithm [24]. Aligned reads were filtered with a mapping quality greater than 10 (MAPQ >10) and PCR duplicates were removed. Bigwigs were created for visualization in Integrative Genomics Viewer [28].

### Motif Analysis

Fasta files for the regions of interest were produced using BEDTools [26]. Motif analysis of all p53 bound regions was performed using MEME-ChIP [29], The MEME Suite [30]. The motif discovery and enrichment mode were performed in classic mode using Human motifs. The number of motifs discovered was set to 10 with a maximum motif width set to 20. All other parameters were set to default.

### p53 Protein Purification

Untagged, thermostable, N-terminally truncated human p53 (residues 94-393) and its Val158 to F (V157F) and Arg158 to L (R158L) mutants were expressed in BL21-CodonPlus (DE3)-RIL E. coli (Stratagene). 6L of bacteria cell cultures were grown at 37°C until they reached an OD600 equal to 0.5–0.8. Cells were then shifted to 18°C. Protein expression was induced by the addition of 0.2mM isopropyl-β-D-thiogalactoside (IPTG). Cells were harvested by centrifugation (20 minutes at 7500 × *g* and 4°C), resuspended in cell lysis buffer [20 mM bis-tris propane (BTP, pH 6.8), 200mM NaCl, 2mM DTT, and 0.5mM Tris (2-carboxyethyl) phosphine hydrochloride (TCEP)] and homogenized by an Emulsiflex C5 cell disruptor (Avestin). Cell lysates were centrifuged at 35,000 × *g* for 30 minutes at 4°C and cleared cell extracts were loaded onto a HiTrap SP cation exchange column (GE Healthcare) pre-equilibrated with a lysis buffer. The proteins were eluted using a linear gradient of NaCl from 200mM to 1M concentration. Eluted proteins were then loaded onto a HiTrap Q anion exchange column (GE Healthcare) to remove residual nucleic acids, and the protein-containing flow-through was collected. Finally, the pooled protein was concentrated and loaded onto a HiLoad Superdex 200 16/60 size exclusion column (GE Healthcare) and eluted in a buffer containing 150mM NaCl, 20mM BTP (pH 6.8), and 0.5mM TCEP. Final protein concentrations were estimated by measuring absorbance at 280nm.

### Colloidal Blue Staining

Colloidal blue staining was performed with a colloidal blue staining kit (LC6025, Invitrogen) following manufacturer protocol for Novex® Tris-glycine gels. Samples were loaded into the gel in addition to either Benchmark Protein Ladder (110747012, Thermo Fisher Scientific) or Precision Plus Protein Dual Color Standards (#1610374, Bio-Rad) for a ladder. The electrophoresis process was conducted at 100V. The gel was stained for 3 hours with colloidal blue solution and then destained for 8 hours with deionized water before being imaged.

### Surface Plasmon Resonance (SPR)

SPR experiments were performed using a Biacore X100 instrument (GE Healthcare) at 25°C using streptavidin coated sensor chips (Sensor chip SA, Biacore X100; GE Healthcare). Sensor chips were primed with running buffer (20mM BTP (pH 6.8); 200mM NaCl; 50µg/mL BSA; 0.005% Tween-20; 0.5mM TCEP) until resonance units (RUs) on all flow cells were stable. Biotinylated dsDNAs were resuspended at a final concentration of 10nM in running buffer and immobilized on the SA sensor chip by injecting at a flow rate of 10µL/min until RUs reached 250. Each experiment utilized two flow cells; DNA was immobilized on one flow cell, and the other flow cell served as a reference. DNA sequences listed in Supplemental Table 3.

To determine p53 DNA binding affinity constants, p53 protein solutions (1, 10, 25, 50, 100, and 200nM protein concentrations, diluted in running buffer) were delivered to the flow cell with immobilized dsDNA and reference cell at a flow rate of 30µL/min for 300 seconds to measure association, followed by dissociation where only running buffer was flowed at 30µL/min for 240 seconds. Between experiments, the sensor chip surface was regenerated by two 120-second injections of running buffer containing 500mM NaCl at 30µL/min to remove any remaining p53 protein from the dsDNA.

Kinetic parameters (association (ka), dissociation (kd), and affinity (KD)) were obtained using BIAevaluation software 2.1 (GE Healthcare). First, RUs collected for the flow cell containing immobilized dsDNA were subtracted by RUs obtained from the reference cell. Then, sensorgrams were globally fitted for all p53 protein concentrations to the Langmuir binding model of simple 1:1 bimolecular interaction. Goodness of fit was evaluated based on the χ2 value and visual inspection.

### Statistical Analysis

Data are expressed as mean±standard deviation. All statistical analysis was performed using GraphPad Prism 10 Version 10.3.1 (464) unless stated otherwise. The data are a representation of three independent biological replicates unless stated otherwise.

## Results

### The V157F p53 mutant binds DNA in endogenously expressing cells

The V157F p53 mutant regulates a novel transcriptome [31, 32]. However, the mechanism behind this regulation is unknown. To determine whether the V157F p53 mutant binds DNA to perform gain-of-function activities, proximity ligation assay (PLA) was performed. The human lung adenocarcinoma line H2087 is homozygous for the V157F p53 mutation. In these cells, significantly increased proximity between p53 and DNA was observed (Figure 1A, Supplemental Figure 1). In fact, quantification of these images demonstrated that the H2087 experimental condition (p53 + Biotin) had significantly higher PLA signal compared to the three control conditions (Figure 1B). These results suggest that the V157F p53 mutant is in close proximity to DNA and may be bound to DNA.

**Figure 1:**
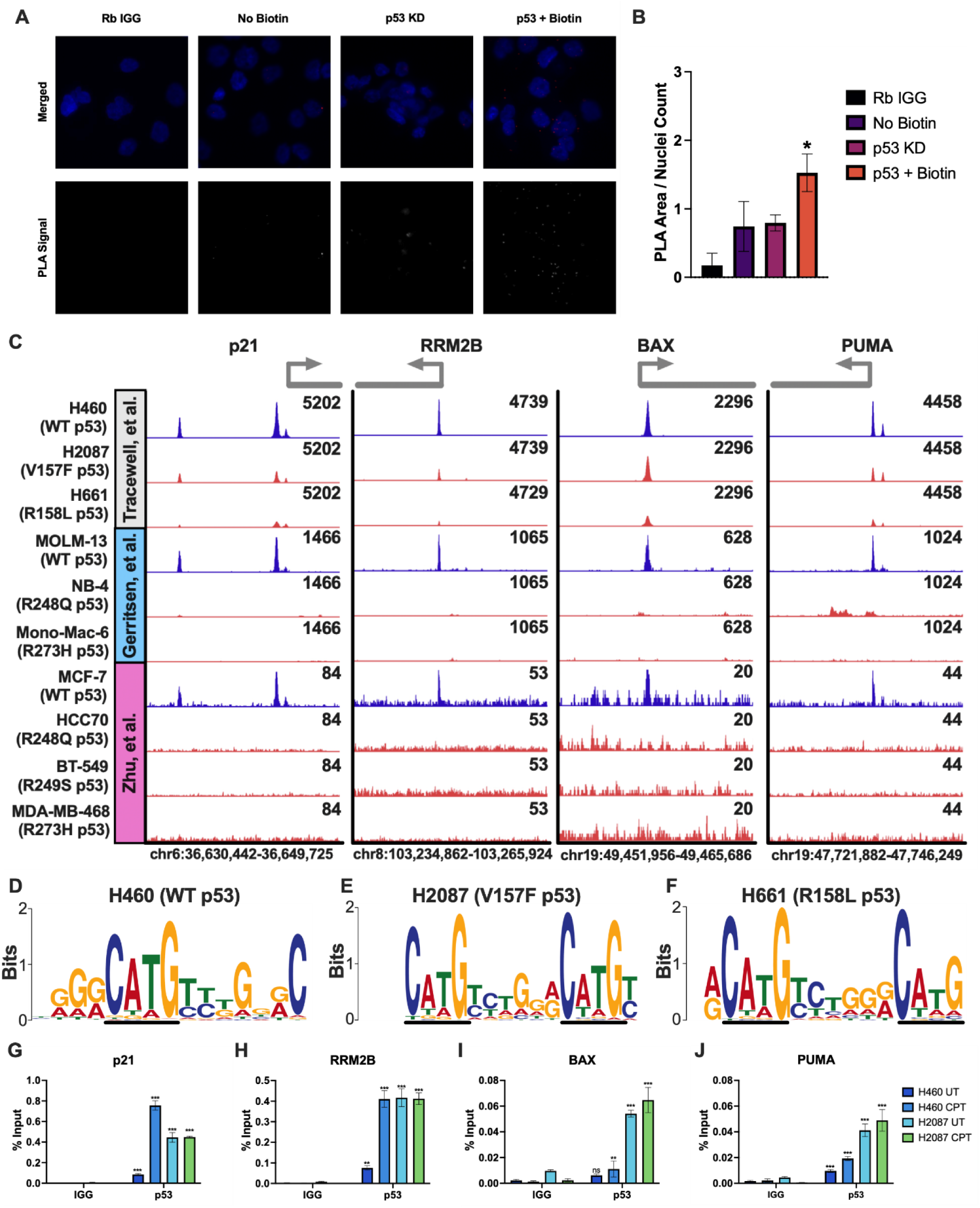
V157F and R158L p53 mutants bind at WT p53 target genes in endogenous cells. **A,** Proximity ligation assay (PLA) was performed on H2087 (V157F p53) cells. Samples included the experimental condition (p53 + Biotin), rabbit IgG in place of the biotin antibody (Rb IgG), a sample with no biotin azide added to the click-it reaction to mark the DNA (no Biotin), and a sample where p53 was depleted from the cells with shRNA (p53 KD). Images shown are of merged DAPI (blue) and PLA signal (red) channels, and PLA signal alone (white). **B,** PLA images containing 100-300 cells per image were quantified by the area of the PLA signal over the nuclei count in the image. p53 + Biotin was significantly different from all three controls by one-way ANOVA, * p-value <0.05. **C,** ChIP-seq was performed on H460 (WT p53) and H2087 (V157F p53) cells treated with CPT (2μM) for 6 hours (Tracewell, et al.). Data from two other publicly available datasets were used (GSE238181 [35] and GSE59176 [36]) that feature cell lines with both WT p53 and different p53 mutants. Signal intensity of p53 binding at representative p53 target genes p21, PLK3, BAX, and PUMA visualized in Integrative Genomics Viewer [28]. Peak scaling was grouped separately for each experiment. WT p53 tracks are blue, and p53 mutant tracks are red/orange. **D-F,** Enriched query motifs at p53 (WT, V157F, or R158L) bound regions identified by MEME-ChIP [29, 30]. The black line highlights the conserved p53 consensus core sequence “C(A/T)(A/T)G”. **G-J,** ChIP-qPCR was performed on H2087 and H460 (WT p53) cells. The ChIP samples were analyzed by qPCR with primers designed at WT p53 binding sites of p53 target genes: p21, RRM2B, BAX, and PUMA. A two-way ANOVA was performed to determine significant differences in binding between the p53 ChIP sample and the corresponding IgG control, *** p-value <0.001, ** p-value <0.01, ns: non-significant.

Lung cancer cells accumulate p53 mutations in a small cluster of amino acids (V157, R158, A159) that are not commonly mutated in other forms of cancer. Given their physical proximity, it was of interest to assess whether other missense mutations at this cluster have similar biological effects. V157F and R158L missense mutations of p53 are the most common alterations in this lung cancer cluster, and we previously showed that these two missense mutations have a similar impact on oxidative stress response [32, 33]. ChIP-seq was performed to determine where V157F and R158L mutant p53 may be bound in the genome. H2087 (V157F p53), H661 (homozygous for the R158L p53 mutant allele), and H460 (WT p53) cells were treated with camptothecin (CPT). CPT binds and inhibits topoisomerase I, causing DNA damage during S-phase of the cell cycle and leading to activation of WT p53 [34]. V157F p53 (H2087) and R158L p53 (H661) were bound at many of the same sites as WT p53 (H460), including canonical WT p53 target genes for both cell cycle arrest genes, such as p21 and RRM2B, and pro-apoptotic genes, such as BAX and PUMA (Figure 1C). In H460 (WT p53) cells, 282 peaks were identified at promoters. Of these 282 promoters bound by WT p53, 241 were bound by V157F (H2087) and 223 were bound by R158L (H661). While V157F and R158L p53 mutants are bound at the same location as WT p53, they have less signal intensity at these sites compared to WT p53.

Two publicly available ChIP-seq datasets (GSE238181 [35] and GSE59176 [36]) that include data from WT p53 cell lines (MOLM-13 and MCF7) and mutant p53 cell lines (NB-4:R248Q, Mono-Mac-6:R273H, HCC70:R248Q, BT549:R249S, MDA-MB-468:R273H) were utilized to demonstrate that mutant p53 variants are not commonly bound at WT p53 loci. Of these publicly available ChIP-seq data, WT p53 was bound at the same loci as the WT p53 (H460), V157F p53 (H2087), and R158L p53 (H661) (Figure 1C). The five different p53 mutants from the publicly available ChIP-seq data had no binding at WT p53 target genes (Figure 1C). The results from a motif analysis for WT p53 (H460), V157F p53 (H2087), and R158L p53 (H661) are shown, and all three have a strikingly similar positional weight matrix for the p53 canonical binding sequence (Figure 1D-F). These results suggest that the V157F and R158L p53 mutants bind at WT p53 target genes, which is unique for p53 mutants.

To verify the finding that V157F p53 mutant binds at WT p53 target genes from the ChIP-seq analysis, ChIP-qPCR was also performed on H2087 (V157F) and H460 (WT) cells. The binding sites of WT p53 at target genes p21, RRM2B, BAX, and PUMA were used to verify that mutant p53 binds to the same genomic region as WT p53 (Figure 1G-J). V157F p53 (H2087) was bound at canonical WT p53 target genes at levels similar to WT p53 with CPT treatment (Figure 1G-J). CPT treatment increased WT p53 (H460) binding at canonical WT p53 target genes as expected, and V157F p53 (H2087) was bound at similar levels with or without CPT stimulus (Figure 1G-J). These results suggest that the V157F p53 mutant is able to bind at canonical WT p53 target genes, which is an unprecedented finding for a p53 mutant.

### V157F and R158L p53 mutants bind DNA similar to WT p53 in vitro

To further support the PLA and ChIP data, the interaction kinetics between p53 and different DNA sequences were studied using a surface plasmon resonance (SPR) approach. A thermostable, N-truncated version of p53 was used for these experiments, previously shown to retain the same DNA binding ability as WT p53 protein (Figure 2A) [18, 37, 38]. WT, V157F, and R158L versions of p53 were purified for use with SPR (Figure 2B). Salt concentrations of 200mM and 275mM were utilized as they have been previously shown in published works to greatly decrease non-specific binding [18]. WT p53 target genes p21 and BAX, along with a scrambled DNA sequence as a control, were used to test the binding affinity for the p53 variants at salt concentrations of 200mM and 275mM, and protein concentrations ranging from 1nM to 200nM (Figure 2C, Supplemental Figure 2). The interaction between the p21 response element sequence and either of the p53 mutants resulted in a slightly lower maximum resonance unit (RU) value than that of the interaction between the p21 response element and WT p53 (Figure 2D). There was no difference in binding interaction between the three p53 variants and the BAX response element (Figure 2E). All three p53 variants did not have a strong binding affinity for the scrambled sequence as expected (Figure 2F). The DNA binding affinity for each of the p53 variants was determined and expressed through the dissociation constant KD (Table 1). The KD for p53 mutants at WT p53 target gene response elements are strikingly similar to the KD of WT p53.

**Figure 2:**
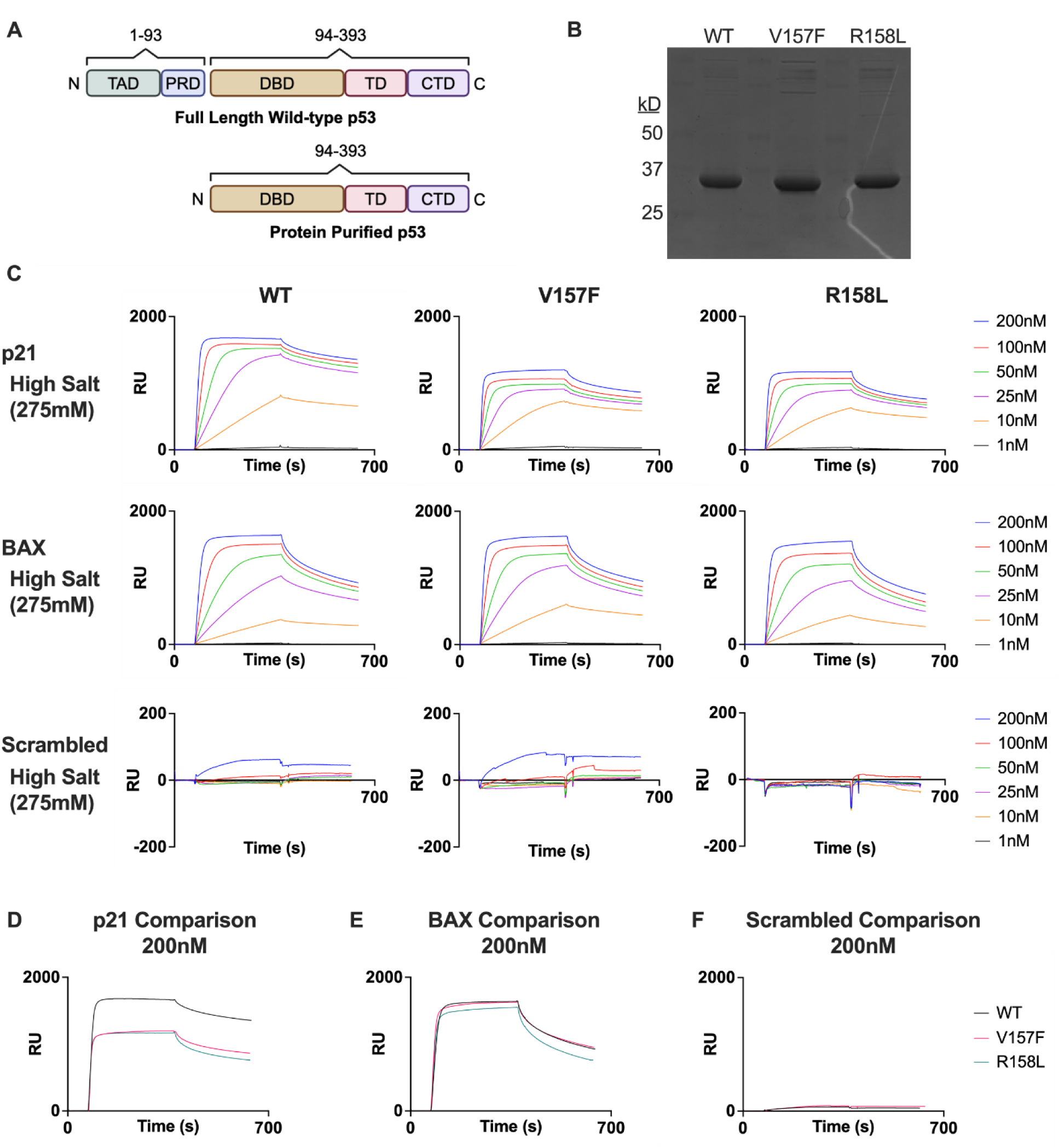
V157F and R158L p53 mutants bind to p53 target genes similar to WT p53. **A,** Schematic of full-length WT p53 and protein purified ΔN-p53 construct used for subsequent SPR experiments. **B,** Coomassie Brilliant Blue staining for protein purified ΔN-p53 constructs of WT, V157F, and R158L p53. **C,** SPR binding data in resonance units (RU) for WT, V157F, and R158L p53. 1nM-200nM protein concentrations were used at a salt concentration of 275mM. Binding was checked using p21 and BAX response elements, as well as a scrambled control DNA sequence. **D-F,** Comparisons between WT and mutant p53 displaying the 200nM protein concentration at 275mM salt concentration for p21 (D), BAX (E), and scrambled (F).

**Table 1:** KD values derived from SPR data for WT p53 and mutant p53 binding interactions with different DNA sequences at different salt concentrations.

|  | WT p53 |  | V157F p53 |  | R158L p53 |  |
| --- | --- | --- | --- | --- | --- | --- |
| Salt Concentration: | 200mM | 275mM | 200mM | 275mM | 200mM | 275mM |
|  | KD (nM) |  | KD (nM) |  | KD (nM) |  |
| p21 | 1.35 | 3.09 | 1.48 | 2.68 | 1.74 | 3.99 |
| BAX | 1.55 | 11.90 | 1.62 | 7.30 | 2.36 | 11.20 |
| Scrambled | 73.70 | 108.00 | 681.00 | 140.00 | 127.00 | 9710.00 |

The binding interactions between the three p53 variants and WT p53 target genes AEN, PLK3, and PUMA were also tested, and the data suggests that the p53 mutants bind similarly to WT p53 at these genes (Supplemental Figure 3A-B, Supplemental Table 4). This is evidence that the mutant protein can bind WT p53 target gene sequences but raises the question as to whether they can form tetramers. This was tested by using a crosslinking agent, glutaraldehyde, and performing gel electrophoresis to determine whether tetramers are forming for each purified p53 protein. The WT, V157F, and R158L p53 proteins can form dimers and tetramers, indicated by the bands at 2x and 4x the purified protein molecular weight upon addition of glutaraldehyde (Supplemental Figure 4). These results suggest that V157F and R158L p53 mutants can tetramerize and interact with WT p53 response elements similarly to WT p53.

### V157F and R158L p53 mutants bind DNA when expressed in p53-null lung cancer cells

The results suggest that endogenous V157F p53 (H2087) and R158L (H661) are able to bind to DNA. However, to determine whether this observation was cell type-specific property, we utilized a H1299 tetracycline (TET)-inducible expression system to compare the DNA-binding affinity of mutant and WT p53 in a p53-null background. The benefits of this system allow for the protein to be expressed at similar levels and in the same genetic background, thus removing most genomic variability for comparing binding affinities. A representative immunoblot of whole-cell lysate was used to confirm TET-induction of p53 in the H1299 TET-system (Figure 3A). Utilizing ChIP-qPCR, the binding sites of WT p53 at target genes p21 and BAX were used to verify that mutant p53 binds to the same genomic region as WT p53 (Figure 3B). All conditions were found to bind DNA significantly over the IgG control for p21 (Figure 3B). WT, V157F, and R158L were all found to bind significantly over IgG control only in the CPT-treated conditions for BAX (Figure 3B). These results suggest that the V157F and R158L p53 mutants are able to bind at some degree to WT p53 target genes in the H1299 TET-system.

**Figure 3:**
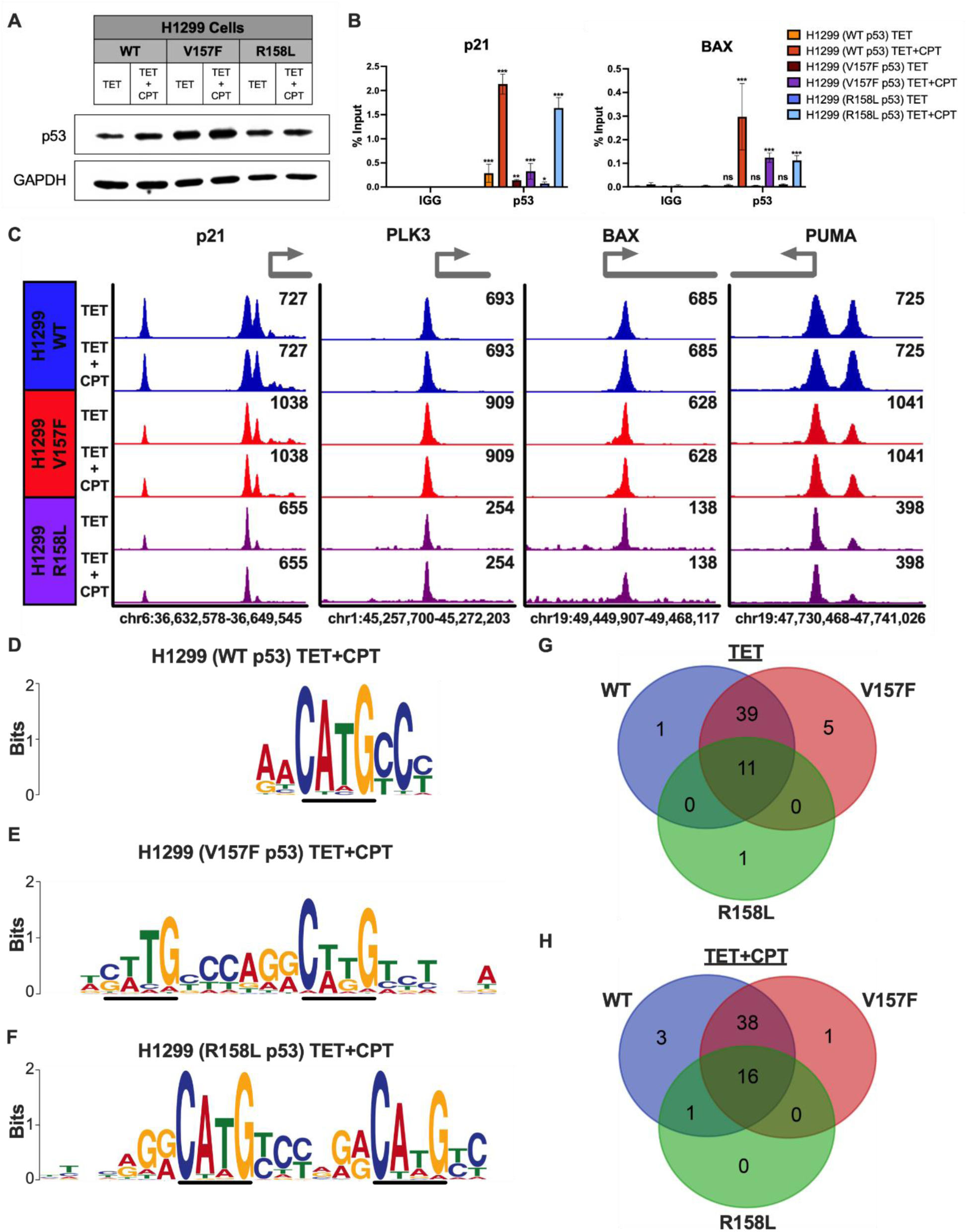
V157F and R158L p53 mutants bind at WT p53 target genes in TET-inducible system. **A,** Representative immunoblot of whole cell lysates collected from samples used for subsequent experiments. The samples consisted of H1299 cells with TET-inducible plasmids for WT, V157F, or R158L p53. Cells were treated with TET alone (1μg/mL) or TET (1μg/mL) followed by CPT (2μM) for an additional 6 hours. The samples had successful induction of p53, and GAPDH was used as a loading control. **B,** ChIP-qPCR was performed on H1299 cells with TET-inducible plasmids for WT, V157F, or R158L p53 treated with TET alone (1μM) and treated with CPT (2μM) for 6 hours or left untreated. The ChIP samples were analyzed by qPCR with primers designed at WT p53 binding sites of p53 target genes: p21 and BAX. Data values given in percentage input. Significance is shown comparing the p53 ChIP sample to the corresponding IgG control, Two-way ANOVA, *** p <0.001, ** p <0.01, * p <0.05, ns: non-significant. **C,** ChIP-seq was performed on H1299 cells containing a TET-inducible plasmid for WT, V157F, or R158L p53 that were treated with TET (1μM) alone for 22 hours or TET (1μM) for 16 hours and then CPT (2μM) for 6 hours. Signal intensity of p53 binding at representative p53 target genes p21, PLK3, BAX, and PUMA visualized in Integrative Genomics Viewer [28]. Peak scaling was grouped separately for each condition; WT p53 (blue), V157F p53 (red), and R158L p53 (purple). **D-F,** Enriched motifs at p53 (WT, V157F, or R158L) bound regions in the TET+CPT-treated condition identified by MEME-ChIP [29, 30]. The black line highlights the conserved p53 consensus core sequence “C(A/T)(A/T)G”. **G-H,** Venn diagrams displaying the number of genes in the KEGG p53 signaling pathway (hsa04115) [39–41] that are bound in WT, V157F, and R158L p53 in both TET-treated and TET+CPT-treated conditions via ChIP-seq.

To further investigate whether mutant p53 can bind DNA at the same areas as WT p53, ChIP-seq was performed on samples containing either TET-inducible WT p53, V157F p53, or R158L p53 induced with TET, and some conditions were additionally treated with CPT to induce DNA damage for robust p53 activity. ChIP-seq peaks for WT p53, V157F p53, and R158L p53 are shown with and without CPT treatment at WT p53 target genes p21, PLK3, BAX, and PUMA (Figure 3C). V157F p53 is bound at representative WT p53 target genes similarly to WT p53, and R158L p53 is clearly bound at WT p53 target genes but to a lesser extent than WT p53, indicated by a lower signal intensity (Figure 3C). The motif analysis for the CPT-treated conditions indicated that WT p53, V157F p53, and R158L p53 all have a similar binding preference for the p53 canonical binding sequence (Figure 3D-F). The KEGG p53 signaling pathway (hsa04115) was utilized to determine the overlap of WT p53 signaling genes identified as bound between WT p53, V157F p53, and R158L p53 from the ChIP-seq analysis [39–41]. A majority of the genes in the KEGG p53 signaling pathway were identified as bound in the V157F p53 and WT p53 groups in the presence or absence of CPT (Figure 3G-H). R158L p53 was found to be bound at a smaller subset of the p53 signaling genes compared to WT p53 or V157F p53 (Figure 3G-H). These results suggest that V157F p53 and R158L p53 expressed in a TET-inducible system bind to canonical WT p53 target genes at levels comparable to WT p53 under endogenous conditions. These data support that this binding is a biological feature of these lung-enriched p53 mutants rather than a cell type-specific property.

### V157F and R158L p53 mutants fail to induce WT p53 target genes

To assess whether the p53 mutants (V157F or R158L) are able to induce the expression of WT p53 target genes that the mutant protein binds to, RNA-seq was conducted utilizing the H1299 TET-expression system. H1299 cells conditionally expressing either WT, V157F, or R158L p53 were treated with CPT to induce DNA damage for robust p53 activation or left untreated. RNA-seq analysis identified 2286 differentially expressed genes (p < 0.05; FDR < 5%; log_2_FC +/-1.5) in the V157F p53 CPT-treated condition compared to the WT p53 CPT-treated condition (Figure 4A). Additionally, 2797 genes were differentially expressed (p < 0.05; FDR < 5%; log_2_FC +/- 1.5) in the R158L p53 CPT-treated condition compared to the WT p53 CPT-treated condition (Figure 4B). Canonical WT p53 target genes were among both sets of differentially expressed genes. In the mutant p53 conditions, p21, PLK3, BAX, and PUMA were significantly downregulated in both the TET and TET+CPT-treated groups compared to the corresponding WT p53 condition, with the exception of PUMA for the R158L CPT-treated group, which was not significantly different from the WT p53 CPT-treated group (Figure 4C). These data suggest that the V157F or R158L p53 mutants are unable to induce expression of WT p53 target genes to a similar level of WT p53 even though they bind to the same target sequence.

**Figure 4:**
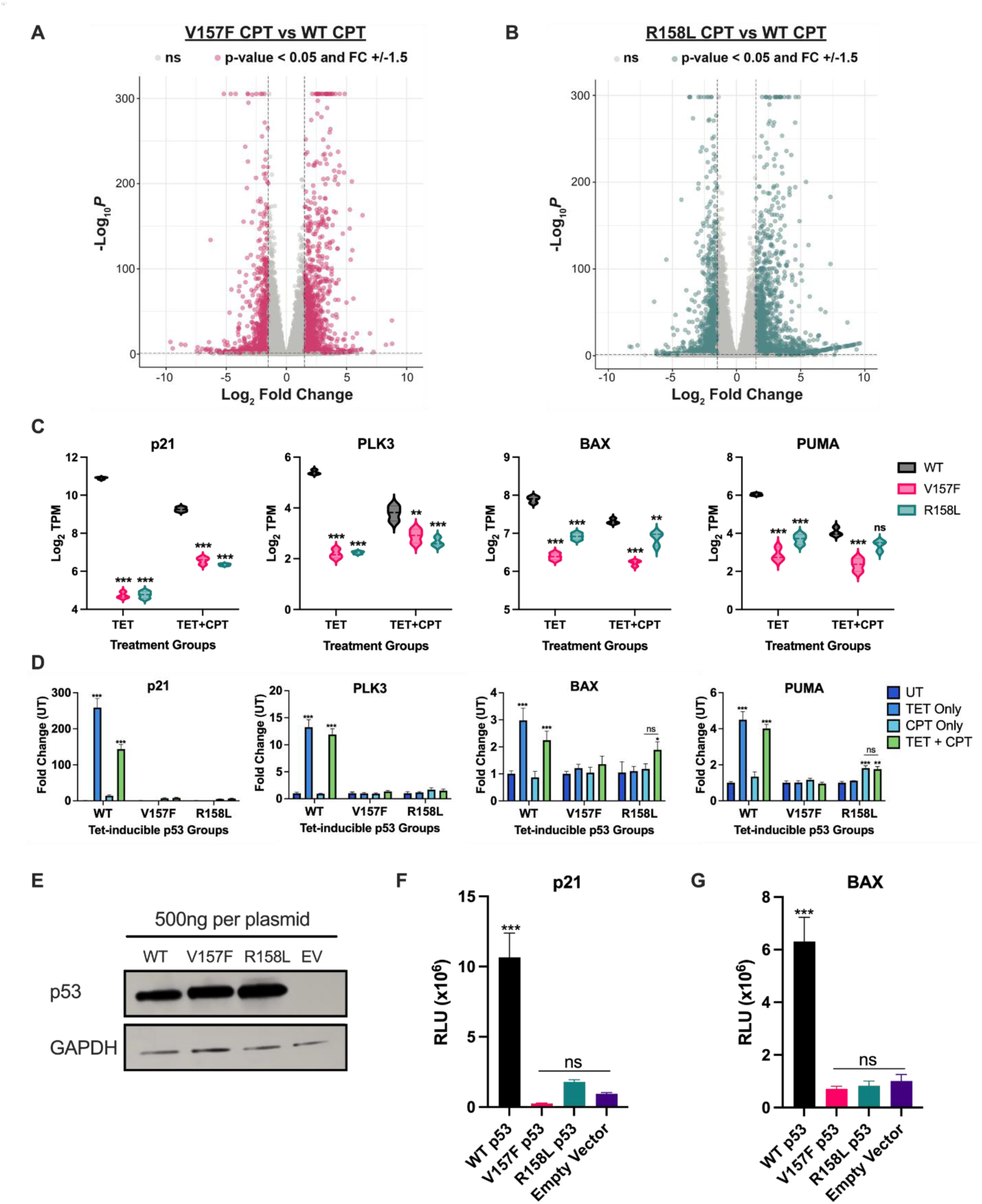
V157F and R158L p53 mutants do not express WT p53 target genes. H1299 cells containing a TET-inducible plasmid for WT, V157F, or R158L p53 were treated with TET (1μM) alone or TET (1μM) and then CPT (2μM) for 24 hours. **A,** Volcano plot of differentially expressed genes in the V157F p53 mutant expressed CPT-treated condition relative to the WT p53 expressed CPT-treated condition determined by DESeq2 [21]. Pink points represent differentially expressed genes (p < 0.05; FDR < 5%; Log_2_Fold Change +/-1.5, n=2286). **B,** Volcano plot of differentially expressed genes in the R158L p53 mutant expressed CPT-treated condition relative to the WT p53 expressed CPT-treated condition determined by DESeq2 [21]. Green points represent differentially expressed genes (p < 0.05; FDR < 5%; Log_2_Fold Change +/-1.5, n=2797). **C,** Violin plots displaying Log_2_TPM of WT p53 target genes p21, PLK3, BAX, and PUMA for TET and TET+CPT treated conditions. Significance is shown for comparison to WT p53 in the respective treatment group, Two-way ANOVA, *** p <0.001, ** p <0.01, ns: non-significant. **D,** RNA was isolated from H1299 cells containing a TET-inducible plasmid for WT, V157F, or R158L p53. The cells were treated with TET only (1μM), CPT only (2μM), TET and CPT, or left untreated. RTq-PCR was performed on the cDNA samples for the WT p53 target genes: p21, PLK3, BAX, and PUMA. Significance is shown for comparison to the UT condition, Two-way ANOVA, *** p <0.001, ns: non-significant. **E,** Whole-cell lysates were collected from H1299 cells 24 hours post-transfection with a luciferase reporter plasmid for p21 or BAX and 500ng of expression plasmid for WT p53, V157F p53, R158L p53, or empty vector (EV). Immunoblot of the whole-cell lysates for p53 and loading control of GAPDH. **F-G,** Lysates from (E) were measured for luminescence in relative light units (RLU) for either p21 (F) or BAX (G). Significance is shown for WT p53 compared to other conditions, One-way ANOVA, *** p <0.001, ns: non-significant.

To confirm the results from the RNA-seq that V157F and R158L p53 mutants are unable to express WT p53 target genes, RT-qPCR was performed utilizing the H1299 TET-expression system. TET expression of WT p53 alone or in combination with CPT treatment induces expression of WT p53 target genes p21, PLK3, BAX, and PUMA (Figure 4D). TET expression of V157F p53 or R158L p53 alone or in combination with CPT treatment does not induce expression of WT p53 target genes p21, PLK3, BAX, and PUMA (Figure 4D). CPT treatment alone without TET expression of p53 has little impact on the expression of WT p53 target genes as expected (Figure 4D). The expression of WT p53 target genes p21, PLK3, BAX, and PUMA was verified in endogenously expressing cell lines H460 (WT p53), H2087 (V157F p53), and H661 (R158L p53) utilizing RT-qPCR. Following CPT treatment, both mutant p53 cell lines had significantly lower expression of WT p53 target genes than WT p53 expressing cells (H460) (Supplemental Figure 5). These data support the RNA-seq findings and suggest that the V157F and R158L p53 mutants are unable to induce functional expression of WT p53 target genes.

A luciferase reporter assay was utilized to further validate that these p53 mutants are unable to induce the expression of representative WT p53 cell cycle control and apoptotic target genes, p21 and BAX, respectively. The same quantity of luciferase reporter plasmid and p53 expression vector encoding either WT p53, V157F p53, R158L p53, or vector alone (EV) was transfected into a H1299 p53-null cell. The cells were lysed, and the lysate was immunoblotted for p53 to show similar p53 levels across the samples (Figure 4E). WT p53 significantly induced the expression of both p21 and BAX (Figure 4F-G). The expression of either V157F or R158L mutant p53 does not significantly induce the expression of either p21 or BAX compared to empty vector control (Figure 4F-G). These results suggest that the V157F and R158L p53 mutants do not induce the expression of canonical WT p53 cell cycle control and apoptotic target genes.

### V157F and R158L p53 mutants are defective for apoptosis induction

We sought to determine whether the observed non-productive binding of these lung-enriched p53 mutants results in a biological consequence. WT p53 has been shown to cause cell cycle arrest and cell death by apoptosis when cells are treated with a DNA-damaging agent [42–44]. CPT was used as the DNA-damaging agent to trigger apoptosis and determine if mutant p53 could induce apoptosis in the same capacity as WT p53. This was first assessed by a cell viability assay in the H1299 TET-expression system (Figure 5A). In CPT-treated conditions, WT p53 expression significantly reduced cell viability compared to either V157F or R158L p53 (Figure 5A).

**Figure 5:**
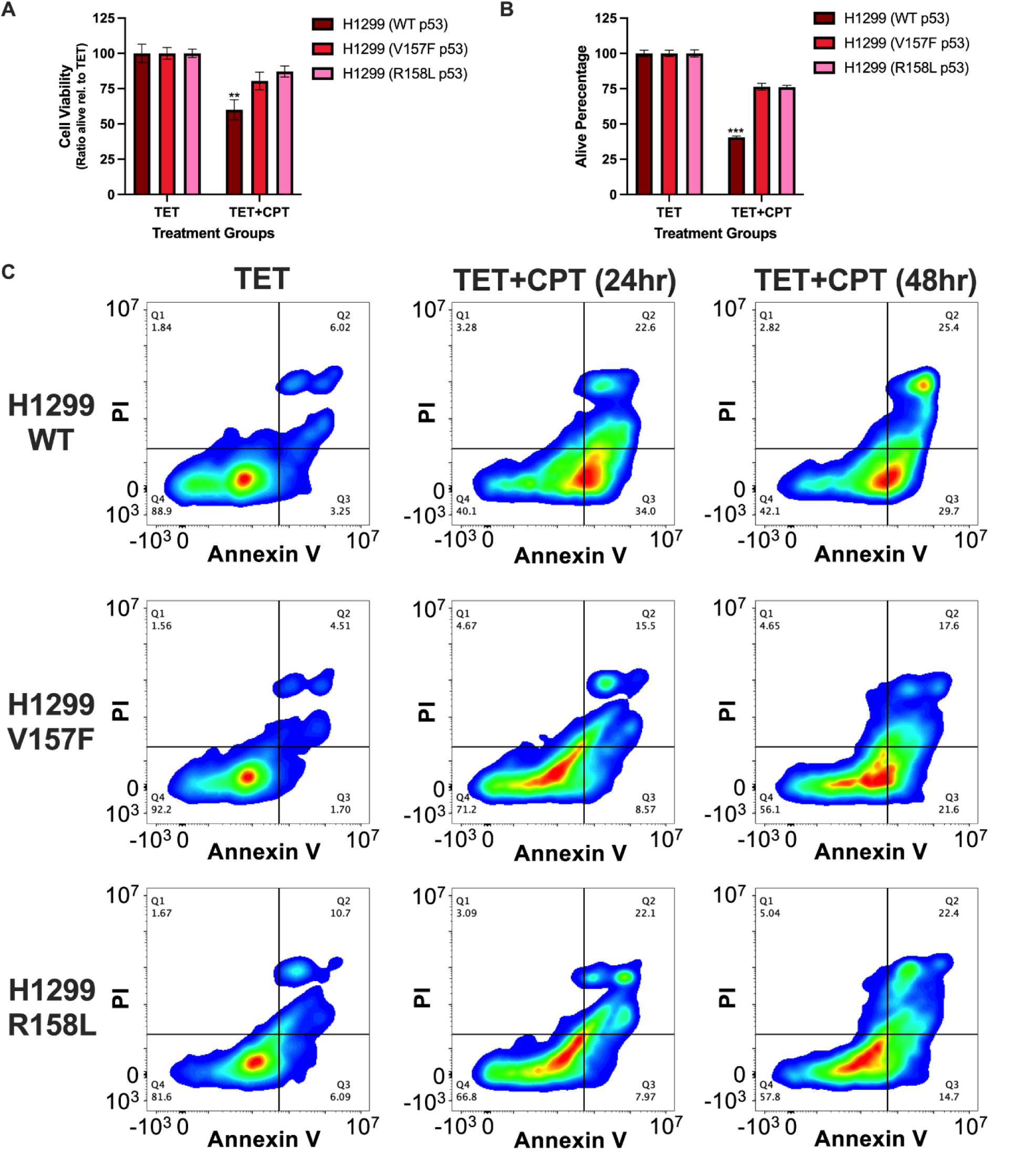
V157F and R158L p53 mutant cells do not induce apoptosis at the same rate as WT p53 cells in response to CPT treatment. For (A-C), H1299 cells containing a TET-inducible plasmid for WT, V157F, or R158L p53 were treated with TET (1μM) alone, or treated with TET (1μM) and CPT (2μM) was additionally added for either 24 or 48 hours. **A,** Cell viability was measured at 48 hours post-addition of CPT. A two-way ANOVA was performed to determine significant differences in cell viability between conditions, significance is shown for comparing TET+CPT conditions, ** p-value <0.01. **B,** Cell quantification was performed on the cells 48 hours post-addition of CPT. A two-way ANOVA was performed to determine significant differences between conditions, significance is shown for comparing TET+CPT conditions, *** p-value <0.001. **C,** Cells were collected and incubated with a staining solution of FITC Annexin V and propidium iodide (PI). The stained cells were analyzed by flow cytometry.

Live cell numbers were also quantified, and under CPT-treated conditions, WT p53 exhibited a significantly lower live cell population compared to either V157F or R158L p53, consistent with productive apoptosis (Figure 5B). Flow cytometry with annexin V and propidium iodide (PI) staining was also used to functionally measure apoptosis. The TET-only cell populations had comparable levels of annexin V and PI staining via flow cytometry among the p53 samples (Figure 5C). CPT-treated cells expressing WT p53 had a rightward shift of the cell population to the second and third quadrants indicating an increase in PI-positive cells, annexin V-positive cells, or double positive cells (Figure 5C). The increase in annexin V-positive cells was drastically higher in CPT-treated cells expressing WT p53 compared to cells expressing V157F or R158L p53 (Figure 5C). These data suggest that V157F and R158L p53 induce apoptosis at a reduced capacity compared to WT p53 following CPT treatment.

To determine whether mutant p53 has an impact on cell cycle arrest, DNA content was analyzed in fixed cells by flow cytometry. The TET-only cell populations had similar percentages of cells in each cycle cell stage: Sub G1, G1, S, and G2/M (Figure 6A-B, Supplemental Table 5). After treatment of CPT for 24 hours, cells expressing WT p53 had a drastic increase in cells with Sub G1 and G1 DNA content compared to cells expressing V157F or R158L p53 (Figure 6A-B, Supplemental Table 5). The cells expressing V157F or R158L p53 with treatment of CPT for 24 hours had a substantial increase in cells with S-phase DNA content and appeared to be in S-phase cell cycle arrest (Figure 6A-B, Supplemental Table 5). After treatment with CPT for 48 hours, cells with V157F or R158L p53 had an increase in cells with Sub G1 and G2/M DNA content compared to the 24-hour CPT-treated conditions (Figure 6A-B, Supplemental Table 5). However, the 48-hour CPT-treated cells expressing WT p53 had the most cells with Sub G1 DNA content, indicating a higher percentage of cells were apoptotic compared to cells with V157F or R158L p53 (Figure 6A-B, Supplemental Table 5). Cell imaging studies further support the flow cytometry data, where cells expressing V157F or R158L p53 undergo cell cycle arrest, and cells expressing WT p53 exhibit increased apoptosis (Supplemental Figure 6A). We next investigated whether mutant p53 exhibited a gain-of-function contributing to increased cell cycle arrest. However, cell cycle arrest was CPT-dependent and not mutant p53-dependent, indicating that CPT induces cell cycle arrest independently of p53 status (Supplemental Figure 6B). In contrast, CPT induced apoptosis specifically in WT p53-expressing cells, whereas mutant p53-expressing cells remained arrested and did not undergo apoptosis to the same extent as WT p53 cells. These data indicate that V157F and R158L p53 exhibit a reduced ability to promote CPT-induced apoptosis compared to WT p53, consistent with a loss-of-function rather than a gain-of-function phenotype.

**Figure 6:**
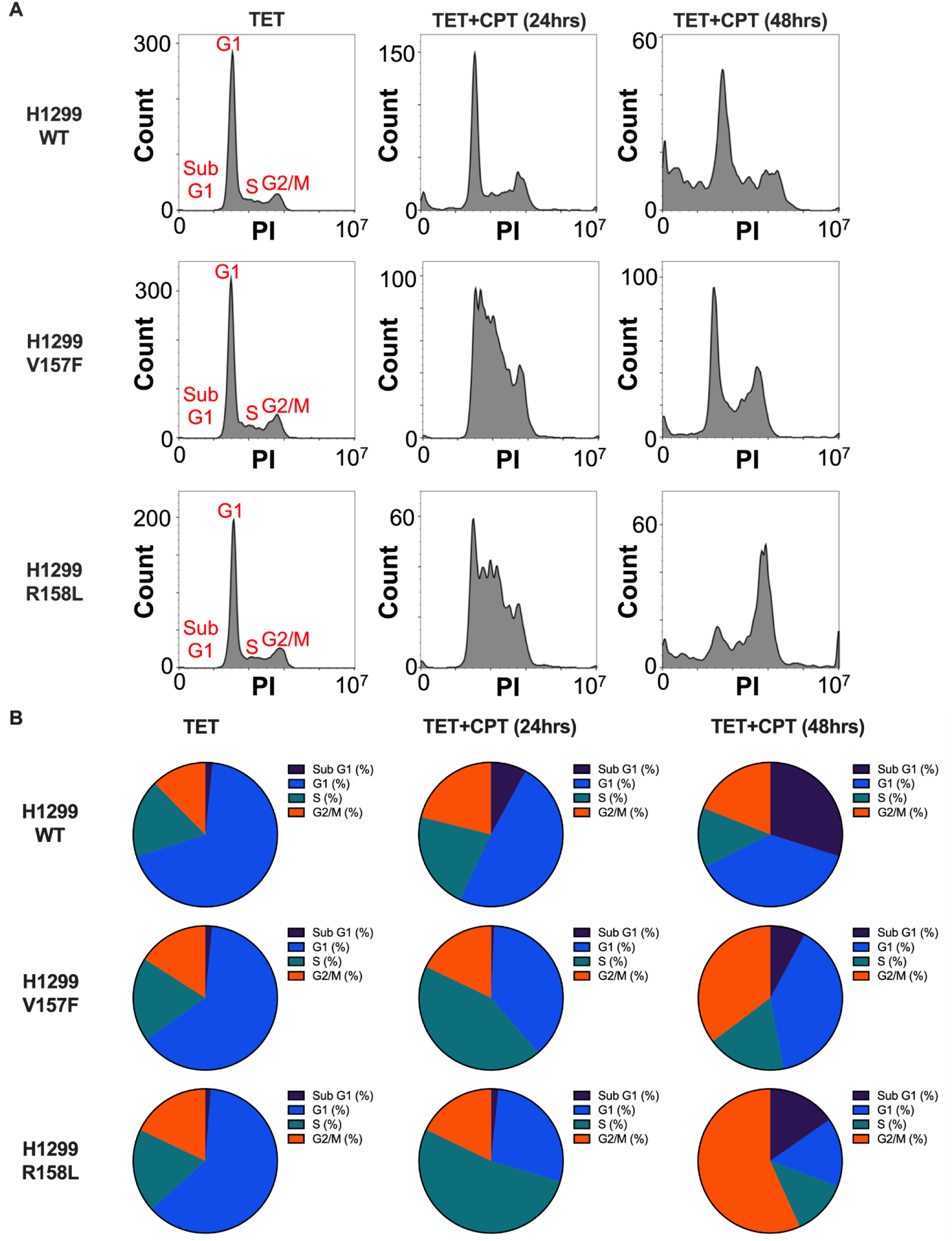
V157F and R158L p53 mutant cells undergo cell cycle arrest, whereas WT p53 cells undergo apoptosis after CPT treatment. **A,** H1299 cells containing a TET-inducible plasmid for WT, V157F, or R158L p53 were treated with TET (1μM) alone, or treated with TET (1μM) and CPT (2μM), which was additionally added for either 24 or 48 hours. Cells were collected, and cell cycle analysis by flow cytometry was performed. Data analysis and histogram creation were done in FlowJo (10.10.0). One-way ANOVA, Two-way ANOVA, p < 0.05. **B,** Pie charts representing the percentage of cells in each stage of the cell cycle (Sub G1, G1, S, and G2/M) are shown from the cell cycle analysis performed in (A).

### V157F and R158L p53 mutants exhibit a dominant-negative phenotype

It was determined that the V157F and R158L p53 mutants participate in non-productive DNA binding at canonical WT p53 target genes. Considering a dominant-negative (DN) phenotype is common amongst p53 mutants, it raises the question of whether these p53 mutants may impact WT p53 function through a DN effect when co-expressed. To assess whether these p53 mutants exert a DN phenotype on WT p53, the H1299 (WT p53) system was utilized. WT p53 was expressed using TET, and either mutant p53 or empty vector were expressed via lipid transfection (Figure 7A). A dual-luciferase reporter assay was performed as a readout for transcription of the p21 and BAX genes. The presence of V157F, R158L, or EV alone showed no significant increase in transcription signal for both p21 and BAX (Figure 7B, 7C). WT p53 induction via TET resulted in a significant increase in transcription of both p21 and BAX in the EV condition as expected (Figure 7B-C). Co-expression of either V157F or R158L p53 mutant with TET-induced WT p53 resulted in a significant reduction in transcription for both p21 and BAX genes (Figure 7B-C). To further support this observation, expression of WT p53 target genes p21, PLK3, BAX, and PUMA were measured by RT-qPCR (Figure 7D-G). Co-expression of either V157F or R158L p53 mutant with TET-induced WT p53 also resulted in a significant reduction in transcription for each WT p53 target gene (Figure 7D-G). These data suggest that V157F and R158L mutant p53 both exert a DN phenotype on WT p53 when co-expressed in a H1299 background.

**Figure 7:**
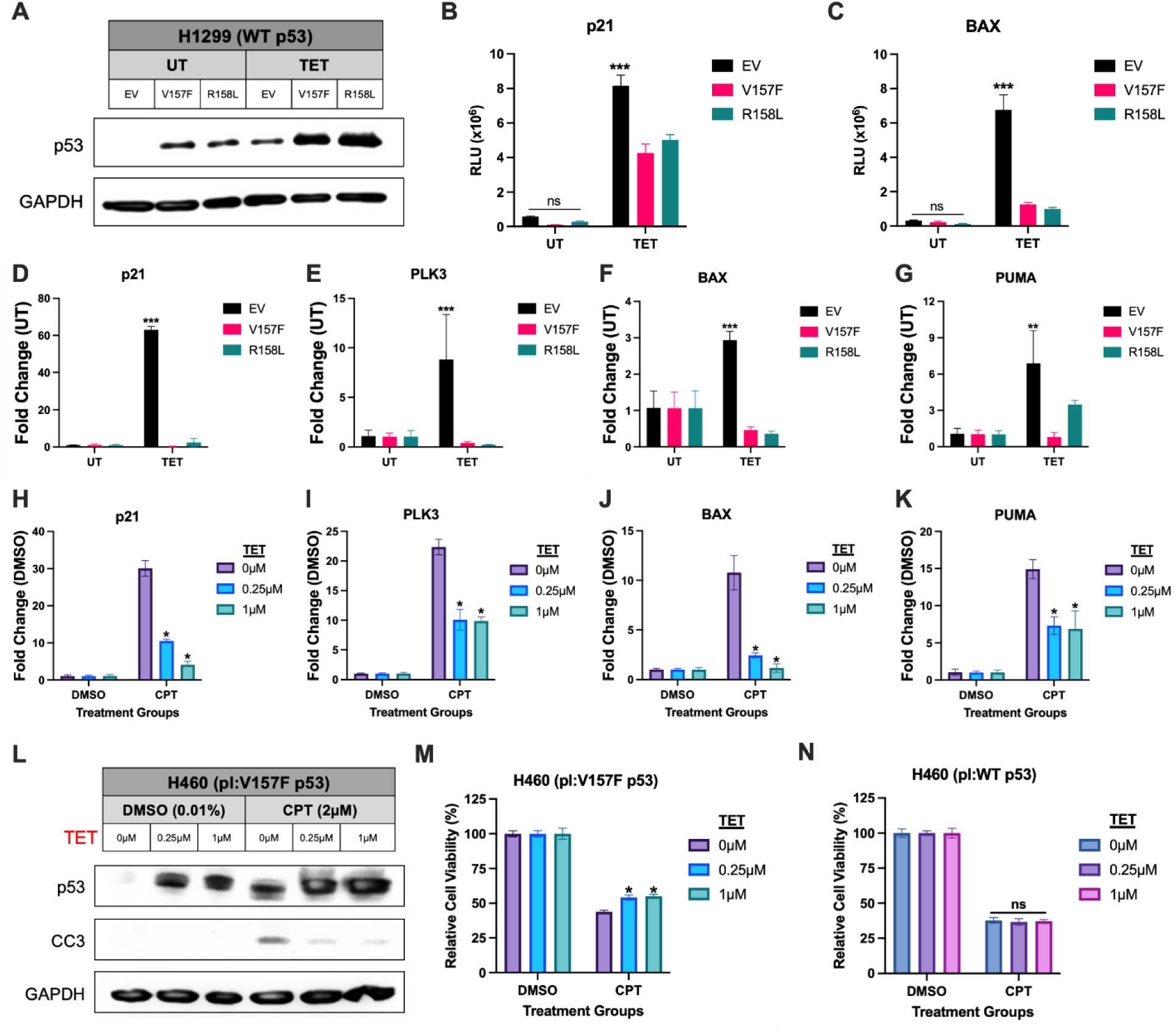
V157F and R158L p53 mutants have a dominant-negative phenotype. **A,** H1299 cells with a TET-inducible plasmid for WT p53 were transfected with a luciferase reporter plasmid for p21 or BAX, and 250ng of expression plasmid for V157F p53, R158L p53, or empty vector (EV). Samples were treated with TET (1μM) to induce expression of WT p53 or left untreated. Immunoblot of the whole-cell lysates for p53 and GAPDH as a loading control. **B-C,** Lysates from (A) were measured for luminescence in relative light units (RLU) for either p21 (B) or BAX (C). Two-way ANOVA was used for statistical analysis, *** p-value <0.001, ns: non-significant. **D-G,** H1299 cells with a TET-inducible plasmid for WT p53 were treated with TET (1μM) to induce WT p53 expression or left untreated. Cells were also transfected with 250ng of expression plasmid for V157F p53, R158L p53, or empty vector (EV). RNA was isolated from the samples, and RTq-PCR was performed on the cDNA samples for the WT p53 target genes: p21 (D), PLK3 (E), BAX (F), and PUMA (G). **H-K,** H460 (WT p53) cells with a TET-inducible plasmid for V157F p53 (pl:V157F p53) were treated with TET (0.25μM or 1μM) or left untreated (0μM). The cells were also treated with CPT (2μM) or equivalent amount of DMSO (0.01%) for 6 hours. RNA was isolated from the cells, and RTq-PCR was performed on the cDNA samples for the WT p53 target genes: p21 (H), PLK3 (I), BAX (J), and PUMA (K). Two-way ANOVA was used for statistical analysis, significance is shown for comparison to 0μM TET + CPT condition, * p-value <0.05. **L,** H460 (WT p53) cells with a TET-inducible plasmid for V157F p53 (pl:V157F p53) were treated with TET (0.25μM or 1μM) or left untreated (0μM). The cells were also treated with CPT (2μM) or equivalent amount of DMSO (0.01%) for 48 hours. Whole cell lysates of the cells were collected and probed for p53, cleaved caspase-3 (CC3), and GAPDH as a loading control. **M,** H460 (WT p53) cells with a TET-inducible plasmid for V157F p53 (pl:V157F p53) were treated with TET (0.25μM or 1μM) or left untreated. The cells were also treated with CPT (2μM) for 48 hours or left untreated. Cell viability was measured at 48 hours post CPT treatment. A two-way ANOVA was performed to determine significant differences in cell viability between conditions, significance is shown for comparison to 0μM TET + CPT condition, * p-value <0.05. **N,** H460 (WT p53) cells with a TET-inducible plasmid for WT p53 (pl:WT p53) were treated with TET (0.25μM or 1μM) or left untreated. The cells were also treated with CPT (2μM) or equivalent amount of DMSO (0.01%) for 48 hours. Cell viability was measured at 48 hours post-addition of CPT. A two-way ANOVA was performed to determine significant differences in cell viability between conditions, significance is shown for comparison to 0μM TET + CPT condition, ns: non-significant.

To assess whether TET-induced mutant p53 would exert a DN phenotype on endogenously expressed WT p53, we utilized the H460 (pl:V157F p53) TET-system that expresses endogenous WT p53 and plasmid-encoded V157F p53 (pl:V157F p53) following TET treatment. WT p53 and subsequent transcription of WT p53 target genes were induced with CPT treatment, and expression of V157F p53 was induced by TET. Expression of WT p53 target genes p21, PLK3, BAX, and PUMA were measured by RT-qPCR (Figure 7H-K). Each WT p53 target gene had significantly increased expression with CPT treatment in the 0μM TET condition with no presence of the V157F p53 mutant (Figure 7H-K). Co-expression of V157F mutant p53 in the 0.25μM and 1μM TET conditions caused a significant reduction in transcription signal of each of the WT p53 target genes (Figure 7H-K). The biological impact of the DN phenotype was further assessed by immunoblotting for the apoptosis marker cleaved caspase-3 (CC3) [45] (Figure 7L). Co-expression of V157F mutant p53 in the 0.25μM and 1μM TET conditions reduced protein levels of CC3 compared to CPT treatment alone (Figure 7L). The biological impact was also assessed using a cell viability assay. Co-expression of V157F mutant p53 with CPT treatment significantly increased cell viability compared to CPT treatment alone in the H460 (pl:V157F p53) TET-system (Figure 7M). In the H460 (pl:WT p53) TET-system, expression of TET-induced WT p53 had no impact on cell viability compared to CPT treatment alone (Figure 7N). Taken together, these data show that the V157F p53 mutant and R158L p53 mutant have DN phenotypes when co-expressed with WT p53.

## Discussion

Loss of p53 function is a near obligatory step in the development and progression of human cancer [46]. Most frequently, p53 inactivation results from individual missense mutations that cause either the generalized misfolding of the protein or the disruption of p53 binding to its canonical genomic sites. Among the mutations that occur most commonly across all cancer types, missense mutations at R175 and Y220 can trigger protein misfolding, and mutations at DNA contact residues, R248 and R273, eliminate sequence-specific DNA interactions [47]. In addition to these ubiquitous hotspot mutations, lung cancer contains an additional hotspot mutation cluster whose activities whose impact is not well-understood. We report here that mutations within this cluster do not cause either p53 misfolding or disruption of its sequence-specific DNA binding capacity. Instead, the function of these p53 mutants is inhibited at a discrete biochemical step subsequent to target gene binding, but prior to transcriptional induction. The identification of non-productive DNA binding by mutant p53 represents a significant advance in our understanding of the biochemical mechanisms responsible for loss of tumor suppressor function in human cancer.

Non-productive DNA binding by the lung-enriched V157F and R158L p53 mutants may relate to these mutants existing in two conformational states, mutant and “WT-like” [48]. The existence of a “WT-like” state for each of these mutants may allow them to bind DNA at WT p53 target genes but not induce gene expression. It is unclear whether the mutant form prevents transactivation of the target gene or if the “WT-like” state is unable to induce gene expression. Therapeutic strategies that force these p53 mutants into a “WT-like” conformation could be sufficient to restore WT function. Restoring WT p53 function is a central goal of many drug development programs targeting mutant p53 [15]. The findings presented here may identify a novel point of intervention for strategies aimed at reconfiguring specific p53 mutants into a “WT-like” state.

Early in the development of most human cancers, incipient tumor cells often harbor one mutant p53 allele while retaining the other WT allele prior to loss of heterozygosity. As p53 functions as a tetramer, the co-expression of mutants and WT alleles result in the formation of inactive heterotetramers [49]. In some cases, even a single monomer carrying a conformational or DNA-binding site mutation can inactivate the tetramer [50, 51]. It remains plausible that a DN phenotype could provide advantages in the earliest stages of tumorigenesis when mutant p53 and wild-type p53 are co-expressed, and help drive the loss of heterozygosity [52, 53]. Furthermore, a DN effect has been shown to be the driving selection force for *TP53* missense mutations in myeloid malignancies [54]. We postulate that the DN phenotype of V157F and R158L may play a minor role in their enrichment in lung cancer.

The discovery that the lung cancer-enriched p53 mutations do not disrupt sequence-specific DNA binding but instead block target gene transactivation raises the possibility that these mutations eliminate the ability of p53 to recruit transcriptional cofactors. Current efforts are focused on understanding whether cofactor recruitment or other events in the transcription cycle are defective in lung cancer cells carrying these p53 mutants.

## Acknowledgements

The authors would like to thank Dr. Jason Hill and BioImaging Core for their help and use of their facilities. The authors thank the Genomic Facility at the Wistar Institute for the Next Generation Illumina Sequencing. The authors thank Dr. Charles Scott and Elizabeth McDuffie for their expertise and use of their equipment. The authors thank Dr. Erik Debler and Dr. Hideharu Hashimoto for assistance with p53 protein purification. The authors thank Dr. Alexander Mazo and Dr. Tyler Fenstermaker for their helpful advice.

## Author Contributions

MAT and SBM designed the project. MAT, HNS, SMB, KMK, TQ, and JEK contributed to experiments. MAT, HNS, and SMB analyzed data. MAT and JAB provided critical advice and tools for the study. MAT and SM wrote the manuscript. All the authors read and approved the manuscript.

## Ethics Statement

N/A

## Data Availability

The RNA-sequencing and ChIP-sequencing data reported in this study are available in GEO: GSE318292 and GSE318294.

