## Supplemental Figures and Tables for "Lung cancer-enriched p53 mutants occupy canonical p53 target genes without activating transcription, revealing a distinct loss-of-function behavior"

| Name | Sequence | Application |
|---|---|---|
| GAPDH FWD | TGGGCTACACTGAGCACCAG | RT-qPCR |
| GAPDH REV | GGGTGTCGCTGTTGAAGTCA | RT-qPCR |
| p21 FWD | TGTCCGTCAGAACCCATGC | RT-qPCR |
| p21 REV | AAAGTCGAAGTTCCATCGCTC | RT-qPCR |
| RRM2B FWD | ATTGGGCCTTGCGATGGATAG | RT-qPCR |
| RRM2B REV | GAGTCCTGGCATAAGACCTCT | RT-qPCR |
| PLK3 FWD | TTTTCGCACCACTTTGAGGAC | RT-qPCR |
| PLK3 REV | GAGGCCAGAAAGGATCTGCC | RT-qPCR |
| BAX FWD | CCCGAGAGGTCTTTTTCCGAG | RT-qPCR |
| BAX REV | CCAGCCCATGATGGTTCTGAT | RT-qPCR |
| PUMA FWD | GACCTCAACGCACAGTACGAG | RT-qPCR |
| PUMA REV | AGGAGTCCCATGATGAGATTGT | RT-qPCR |

**Supplemental Table 1:** List of primers used in RT-qPCR analyses.

| Name | Sequence | Application |
|---|---|---|
| p21 RE FWD | CTCCATCCCTATGCTGCCTG | ChIP |
| p21 RE REV | AGGCAGCCCAAGGACAAAAT | ChIP |
| PLK3 RE FWD | ACCCGAAATTTGCCCCTCAA | ChIP |
| PLK3 RE REV | CTGCCTTGCCAGGTTTAGGA | ChIP |
| RRM2B RE FWD | TGGGCATGTCAGGACAACAA | ChIP |
| RRM2B RE REV | CCAGCCTAGATTGGGGCATC | ChIP |
| BAX RE FWD | TGCGATCTCCAAGCACTGAG | ChIP |
| BAX RE REV | CTTCCGAGATCCCCTGGTTC | ChIP |
| PUMA RE FWD | GAACCCCACACAAACAGGC | ChIP |
| PUMA RE REV | GTGGTGGTGTCGATCCAGAG | ChIP |

**Supplemental Table 2:** List of primer sequences used in ChIP-qPCR analyses.

| Name | Sequence | Application |
|---|---|---|
| p21 FWD | 5BiosG/ATGATGATGAACATGTCCCAACATGTTGATATGATG | SPR |
| p21 REV | CATCATATCAACATGTTGGGACATGTTCATCATCAT | SPR |
| BAX FWD | 5BiosG/ATGATGATGGGCAGGCCCGGGCTTGTCGATATGATG | SPR |
| BAX REV | CATCATATCGACAAGCCCGGGCCTGCCCATCATCAT | SPR |
| Scrambled FWD | 5BiosG/ATGATGATATGAGCACTGTCGTCATAACATATGATG | SPR |
| Scrambled REV | CATCATATGTTATGACGACAGTGCTCATATCATCAT | SPR |
| AEN FWD | 5BiosG/ATGATGATGGGCTTGCCCGGGCATGTGGATATGATG | SPR |
| AEN REV | CATCATATCCACATGCCCGGGCAAGCCCATCATCAT | SPR |
| PLK3 FWD | 5BiosG/ATGATGATTAAACATGCCCGGGCAAAAGCATATGATG | SPR |
| PLK3 REV | CATCATATGCTTTTGCCCGGGCATGTTTAATCATCAT | SPR |
| PUMA FWD | 5BiosG/ATGATGATGGACAAGTCAGGACTTGCAGATATGATG | SPR |
| PUMA REV | CATCATATCTGCAAGTCCTGACTTGTCCATCATCAT | SPR |

**Supplemental Table 3:** List of DNA sequences used in SPR analyses.

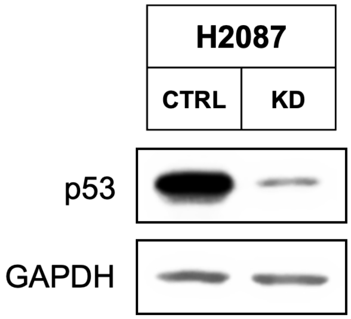


**Supplemental Figure 1:** p53 was successfully depleted in the KD condition.
Immunoblot of GADPH and p53 proteins using whole cell lysates collected from H2087 cells and H2087 p53 KD cells used for the control condition in Figure 1A.

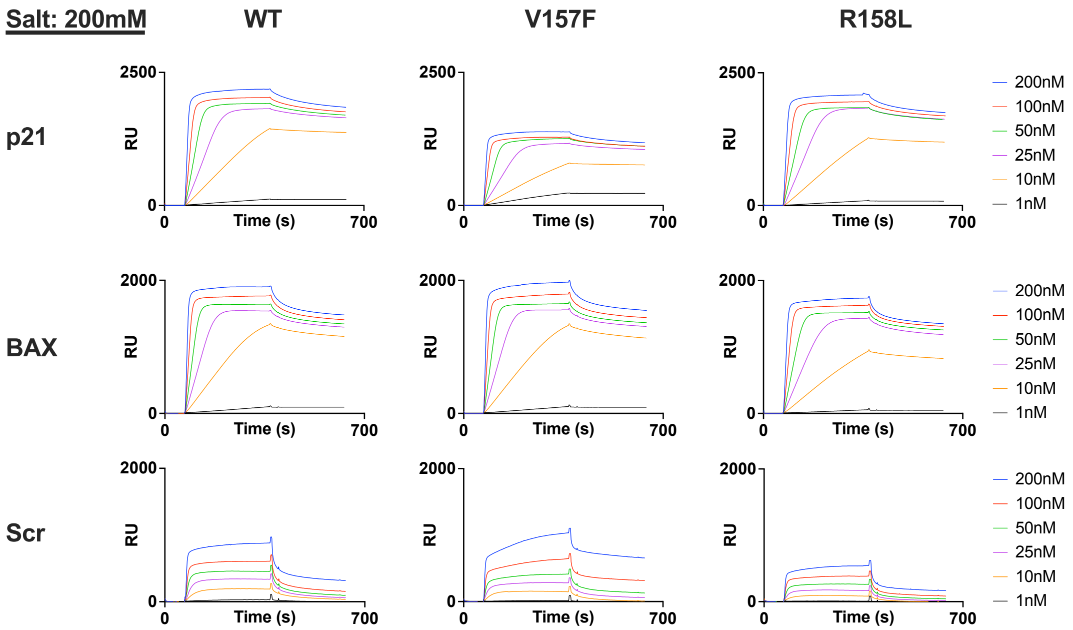


**Supplemental Figure 2:** V157F and R158L p53 mutants bind to p53 target genes p21, BAX, and Scramble similar to WT p53 at 200mM salt by SPR.

SPR binding data in RU for WT, V157F, and R158L p53 at protein concentrations of 1nM-200nM and salt concentration of 200mM. Binding was checked using p21 and BAX response elements and a scrambled control DNA sequence (Scr).

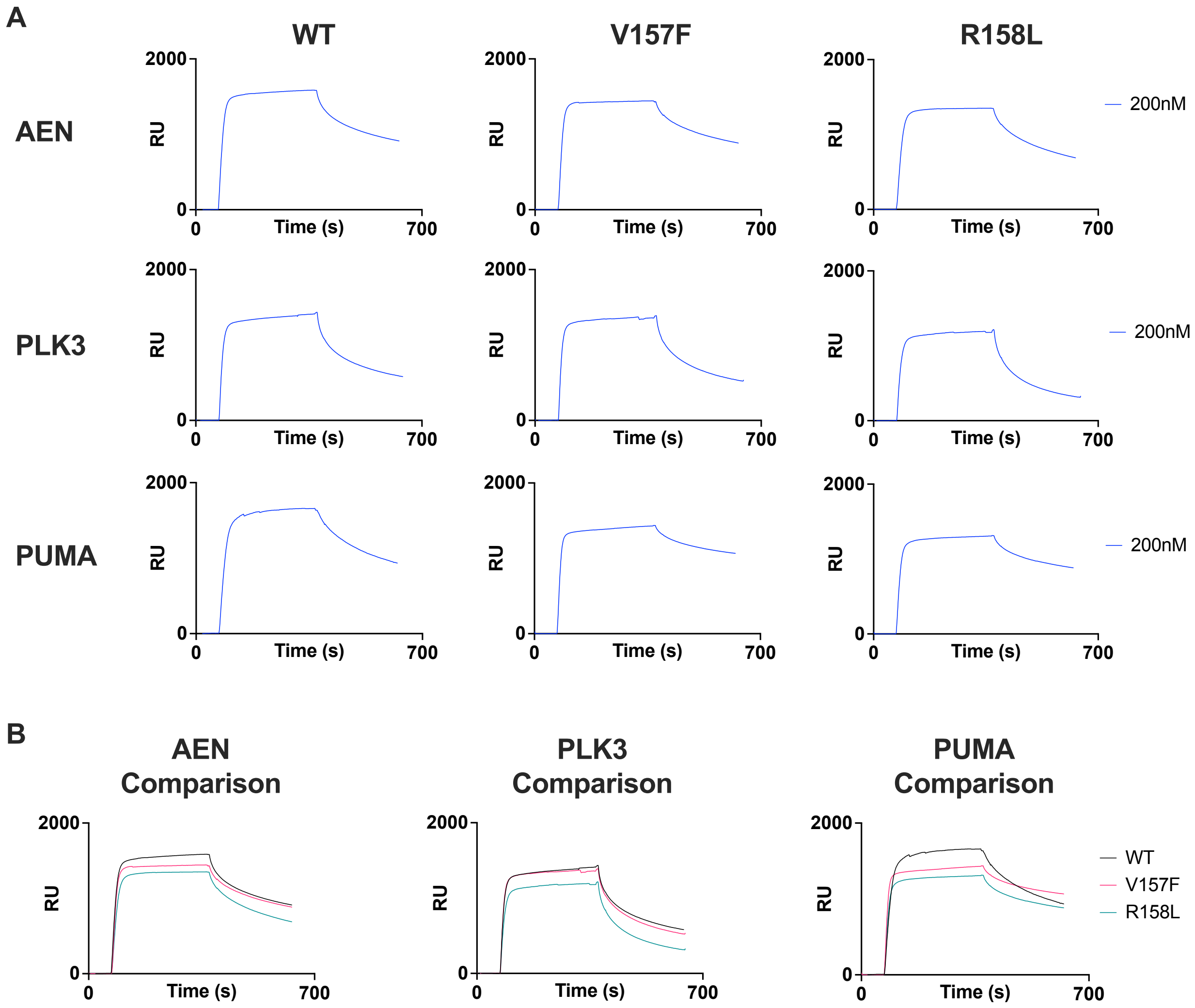


**Supplemental Figure 3:** V157F and R158L p53 mutants bind to p53 target genes AEN, PLK3, and PUMA similar to WT p53 at 275mM salt by SPR.

**A,** SPR binding data in RU for WT, V157F, and R158L p53 at 200nM protein concentration and a salt concentration of 275mM. Binding was checked using AEN, PLK3, and PUMA response elements. **B,** Comparisons between WT and mutant p53 displaying the data from (H) for AEN, PLK3, and PUMA response elements.

| | WT p53 | V157F p53 | R158L p53 |
|---|---|---|---|
| Salt Concentration: | 275mM | 275mM | 275mM |
| | KD (nM) | KD (nM) | KD (nM) |
| AEN | 7.14 | 5.83 | 11.50 |
| PLK3 | 10.10 | 10.50 | 21.90 |
| PUMA | 2.83 | 3.15 | 5.33 |

**Supplemental Table 4:** KD values derived from SPR data for WT p53 and mutant p53 binding interactions with DNA sequences for AEN, PLK3, and PUMA at 275mM salt concentration.

| | WT p53 | | | | | R158L p53 | | | | | V157F p53 | | | |
|---|---|---|---|---|---|---|---|---|---|---|---|---|---|---|
| **Glutaraldehyde %** | 0 | 0.001 | 0.005 | 0.1 | | 0 | 0.001 | 0.005 | 0.1 | | 0 | 0.001 | 0.005 | 0.1 |

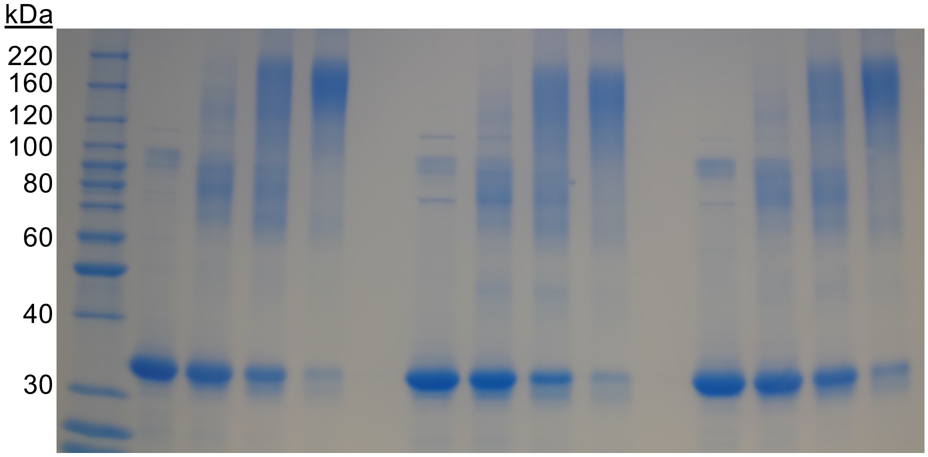


**Supplemental Figure 4:** V157F and R158L p53 mutants are able to form tetramers.
Purified p53 proteins (WT p53, R158L, or V157F p53) were cross-linked with differing concentrations of glutaraldehyde and run on a Coomassie gel.

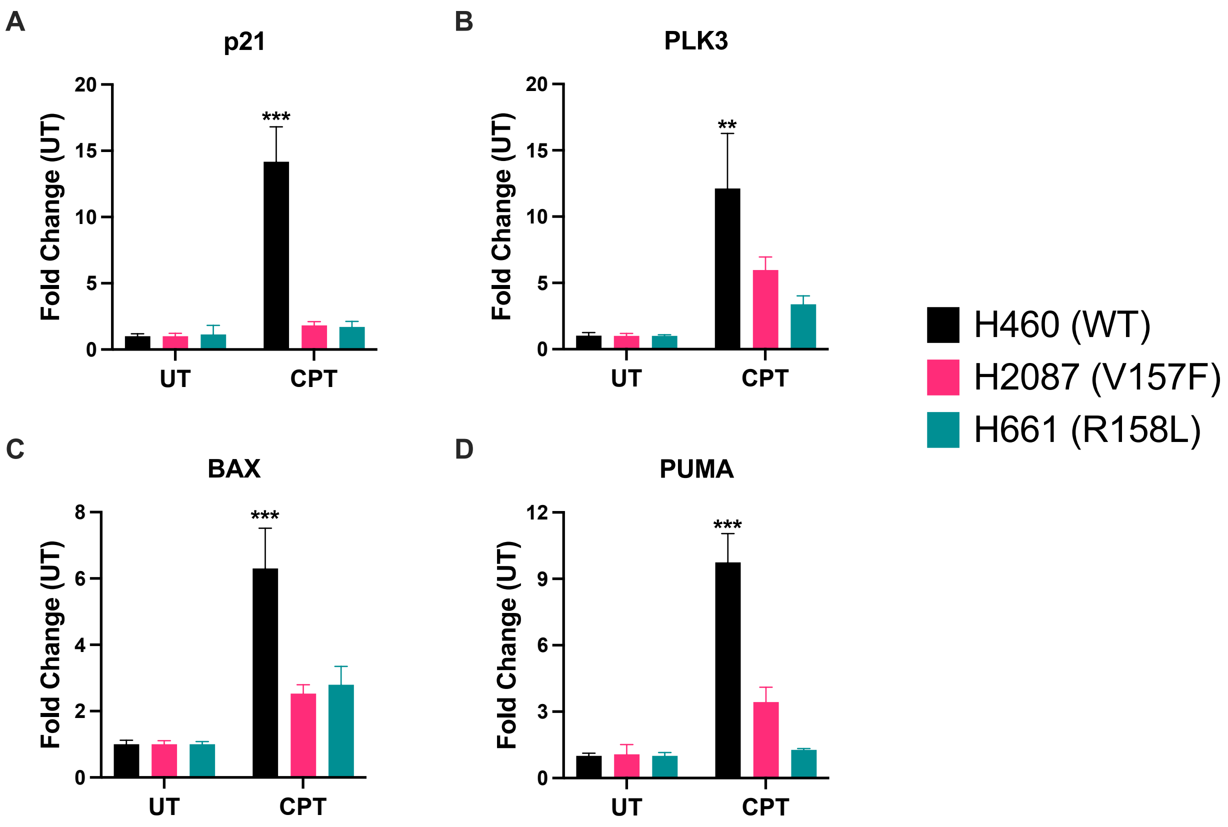


**Supplemental Figure 5:** Endogenously expressing V157F (H2087) and R158L (H661) p53 mutant cells do not induce p53 target gene expression to the same degree as WT p53 (H460) cells in response to CPT treatment.

**A-D,** H460 (WT p53), H2087 (V157F p53), and H661 (R158L p53) cells were treated with CPT (2μM) for 16 hours or left untreated. RNA was collected from the cells, and cDNA was created. RT-qPCR was performed on the samples utilizing primers for p21, PLK3, BAX, and PUMA. Significance is shown for H460 (WT) CPT-treated compared to H2087 (V157F) CPT and H661 (R158L) CPT, Two-way ANOVA, *** $p < 0.001$, ** $p < 0.01$.

| | H1299 (WT p53) | | | H1299 (V157F p53) | | | H1299 (R158L p53) | | |
|---|---|---|---|---|---|---|---|---|---|
| **Cell Cycle** | **TET** | **24hr TET/CPT** | **48hr TET/CPT** | **TET** | **24hr TET/CPT** | **48hr TET/CPT** | **TET** | **24hr TET/CPT** | **48hr TET/CPT** |
| Sub G1 (%) | 1.6 | 7.8 | 29.7 | 1.5 | 0.7 | 7.8 | 1.2 | 1.6 | 14.5 |
| G1 (%) | 67.9 | 47.9 | 37.5 | 62.7 | 37.0 | 38.5 | 61.4 | 27.4 | 14.5 |
| S (%) | 17.4 | 21.3 | 13.3 | 18.5 | 42.3 | 17.4 | 18.5 | 51.5 | 11.9 |
| G2/M (%) | 12.3 | 20.6 | 18.9 | 15.7 | 17.4 | 34.7 | 17.7 | 17.5 | 53.7 |

**Supplemental Table 5:** Percentage of cells in each stage of the cycle in Figure 6A.

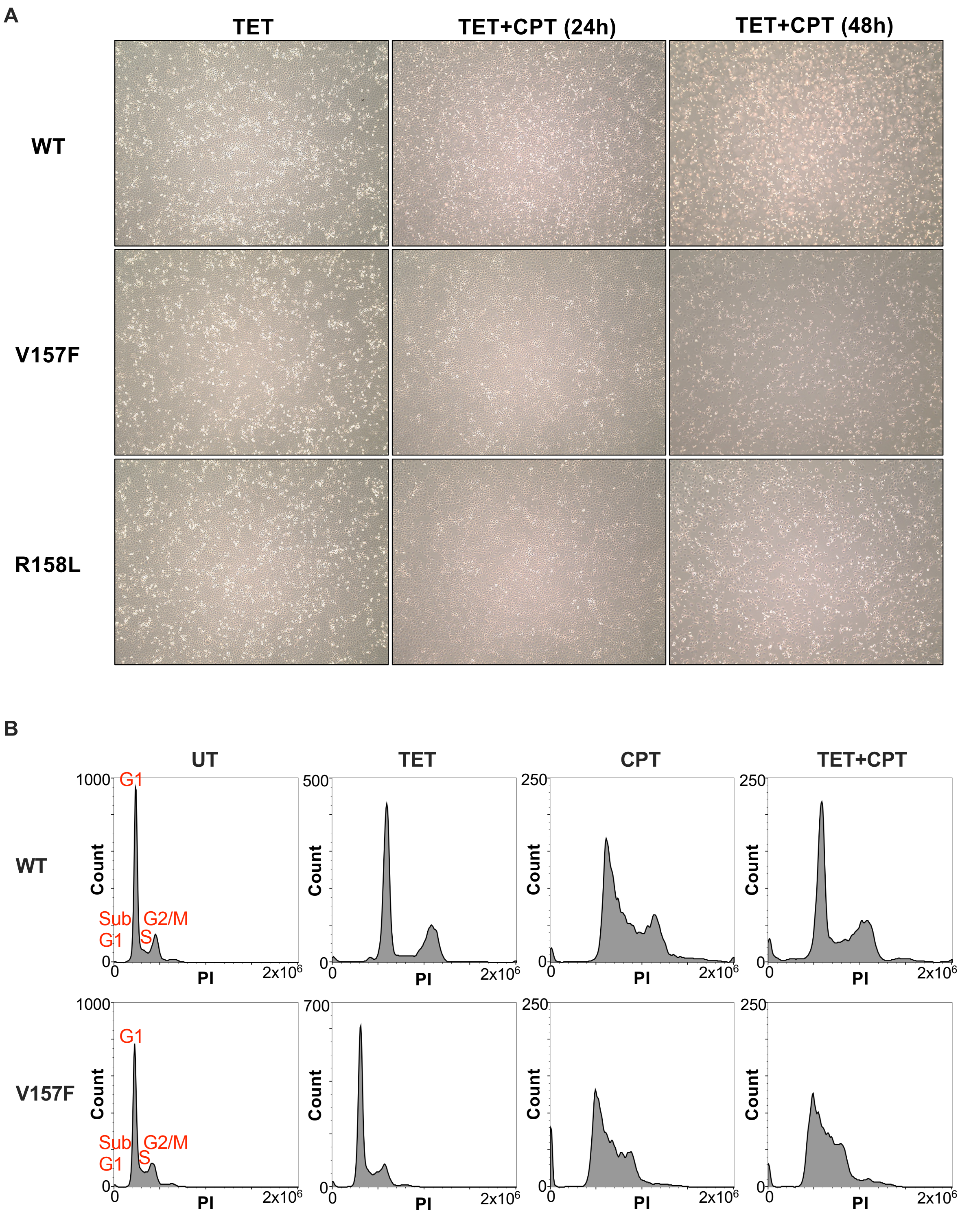


**Supplemental Figure 6:** V157F and R158L p53 mutants have a dominant-negative phenotype.

**A,** Representative brightfield images of H1299 cells used in Figure 6A. Scale bar = 350μm, 4X magnification. **B,** H1299 cells with a TET-inducible plasmid for WT or V157F p53 were treated for 24 hours with TET (1μM) alone, CPT (2μM) alone, TET(1μM) in combination with CPT (2μM), or left untreated. Cells were collected, and cell cycle analysis by flow cytometry was performed. Data analysis and histogram creation were done in FlowJo (10.10.0). Two-way ANOVA, $p < 0.05$.
